# Mcm10 braces converging replicative helicases to pull apart origin DNA

**DOI:** 10.64898/2026.05.15.725099

**Authors:** Giacomo Palm, Agata Butryn, Berta Canal, Thomas Pühringer, Marta H. Gross, John F. X. Diffley, Alessandro Costa

## Abstract

Eukaryotic DNA replication initiation requires controlled, temporally separated steps to preserve genome stability. The MCM replicative helicase is loaded on duplex DNA as an inactive double hexamer, which nucleates the assembly of dimeric Cdc45-MCM-GINS-Pol epsilon (dCMGE) replisomes. Mcm10 splits dCMGE into two, generating divergent replication forks, but the mechanism is unknown. Using ATPase-defective yeast MCM variants that slow origin melting, we captured five reaction intermediates that explain the structural mechanism. Two Mcm10 molecules bridge across the CMGE dimer, bracing two converging helicases. The restrained MCM motors pull apart the two DNA filaments, such that each lagging strand becomes ejected through the Mcm2-5 gate. Our reconstituted structures resemble the double CMG stabilized by metazoan DONSON, pointing to an origin DNA melting mechanism conserved across evolution.

## Introduction

Genome integrity depends on DNA replication occurring exactly once per cell cycle. To achieve this, eukaryotic cells have evolved tightly coupled mechanisms, whereby loading of the replicative helicase onto DNA is permitted only when its activation is suppressed, while the same trigger that activates the helicase also blocks further loading. This process is best understood in yeast, thanks to more than 30 years of genetic and biochemical investigation, culminating in the reconstitution of DNA replication *in vitro* with purified yeast proteins (*1, 2*).

During late mitosis and throughout G1 phase, two copies of the MCM ATPase motor of the replicative helicase are loaded around duplex DNA, forming an inactive double hexamer (DH) held together by dimerising N-termini (*3–7*). Helicase activation occurs at the S phase transition, when two kinases (DDK (*8–10*) and CDK) promote the engagement of a set of assembly factors (Sld2, Sld3, Sld7 and Dpb11) (*11–13*). These recruit three activators (Cdc45, GINS and Pol epsilon) onto the DH, together forming the pre-Initiation Complex (pre-IC), in which the two MCM motors become partially separated (*14*).

In a subsequent step, the assembly factors are released, leaving Cdc45, GINS and Pol epsilon bound to both MCM hexamers, which move farther apart without fully detaching. This state is known as the double Cdc45-MCM-GINS-Pol epsilon (dCMGE) complex (*15*). A further activation step, mediated by Mcm10, splits the dCMGE into two single CMGEs (sCMGEs), which constitute the replication-competent core of the replisome (*16, 17*).

All changes in molecular composition described so far are controlled by the binding, hydrolysis and release of ATP by the MCM motor. During DH formation, for example, each MCM hexamer hydrolyses ATP to translocate away from its loader (ORC-Cdc6) along duplex DNA, such that, by the time the DH is assembled, four ATPase sites are left bound to ADP (*18*). The stepwise recruitment of replisome assembly factors then promotes ADP release (*17*). As a result, within the pre-IC intermediate, ten of the twelve ATPase sites are nucleotide-free (*14*). dCMGE formation requires rebinding of ATP (*15, 17*), while splitting into two sCMGEs is coupled to reactivation of the MCM ATP hydrolysis function (*16, 17*).

These nucleotide-dependent changes alter the way MCM engages DNA, which eventually promotes melting of the double helix. In the DH, the two hexamers are slightly offset, causing duplex DNA to bend as it traverses a narrowing at their interface (*6, 7*). In the pre-IC, partial disengagement of the hexamers allows the double helix to straighten, similar to a compressed spring being released (*14*). ATP binding in the dCMGE then fundamentally changes how MCM grips DNA: within the ATPase tiers at the C-terminal sides of the dimeric complex, the double helix becomes untwisted and base pairing destabilised, while B-form DNA is maintained where it is encircled by the two N-terminal tiers at the homodimerisation core (*15*). By the time the sCMGE is formed, MCM has transitioned from encircling duplex to single-stranded DNA (*16*). This requires ejection of the lagging strand template from the central channel of each hexamer and the crossing of the two separated helicases, so that the strand evicted by one sCMGE becomes the translocation strand of the other, and vice versa (*17, 19*).

Recent reconstitution experiments have revealed that dCMGE splitting and lagging strand ejection are tightly coupled. In fact, when Sld2 is omitted from the activation reaction, dCMGE splitting is defective. The minority of complexes that do reach the sCMGE stage remain bound to duplex DNA, meaning that they failed to eject the lagging strand (*14*).

Although it is established that Mcm10 recruitment underlies both dCMGE splitting and lagging strand ejection, the mechanism remains mysterious (*1, 2, 17*). In fact, Mcm10 contains two established functions, single-stranded DNA binding (*20*) and fork rate stimulation (*21*), but neither is essential in the process of activating an origin of replication (*16*). Mcm10, we reasoned, must be able to reshape the dCMGE, enabling strand ejection in ways we do not understand. Capturing Mcm10 while it activates a dCMGE therefore becomes critical.

Unfortunately, our time-resolved cryo-EM attempts to capture this intermediate so far have failed, suggesting that the kinetics of dCMGE splitting and lagging-strand ejection are faster than we can sample. To circumvent these issues and image Mcm10 while it reshapes a dCMGE, we targeted MCM ATPase mutants that slow the helicase activation reaction. Because MCM is a hetero-hexamer (*22*), mutations can be introduced to inactivate any of the six ATPase sites independently. Arginine-to-alanine (RA) changes in the Arginine Finger of Mcm3, Mcm7 and Mcm4 were shown to support CMG formation (with varying degrees of efficiency), but yield defective recruitment of the single-strand DNA binding protein, Replication Protein A (RPA) (*23*), to varying degrees. We speculated that these defective variants would slow lagging-strand ejection, providing an opportunity to image Mcm10 in the act of activating the replicative helicase. To explore our hypothesis, we used cryo-EM to image lagging strand ejection reactions assembled on an origin of replication with ATPase defective MCM mutants. Our data explain how two converging helicases trapped by Mcm10 achieve DNA melting and reveal the path of lagging strand ejection that promotes replication fork establishment. These results also have implications for bidirectional replication fork establishment in higher eukaryotes.

### An ATPase variant defective for dCMGE splitting

To capture Mcm10-dependent helicase activation intermediates, we used Mcm3, Mcm7 and Mcm4 RA variants (Fig. 1A), each of which loads onto origin DNA as DHs with near-wild-type efficiency (*18*). Sequential purification steps first isolated phospho-DHs, then dCMGE or sCMGE complexes assembled with Mcm10 on linear ARS1 DNA substrates capped at both ends by HpaII methyltransferase (MH) (*14*). Negative stain EM analysis of these preparations revealed strikingly different capacities for dCMGE splitting among the three mutants (Fig. 1B and fig. S1A). Whereas Mcm4RA retained substantial splitting activity (63% of wild type), Mcm7RA and Mcm3RA were progressively more impaired (17.5% and 4%, respectively; wild type = 100%; based on 170,398 averageable particles across three biological replicates).

**Fig. 1.**
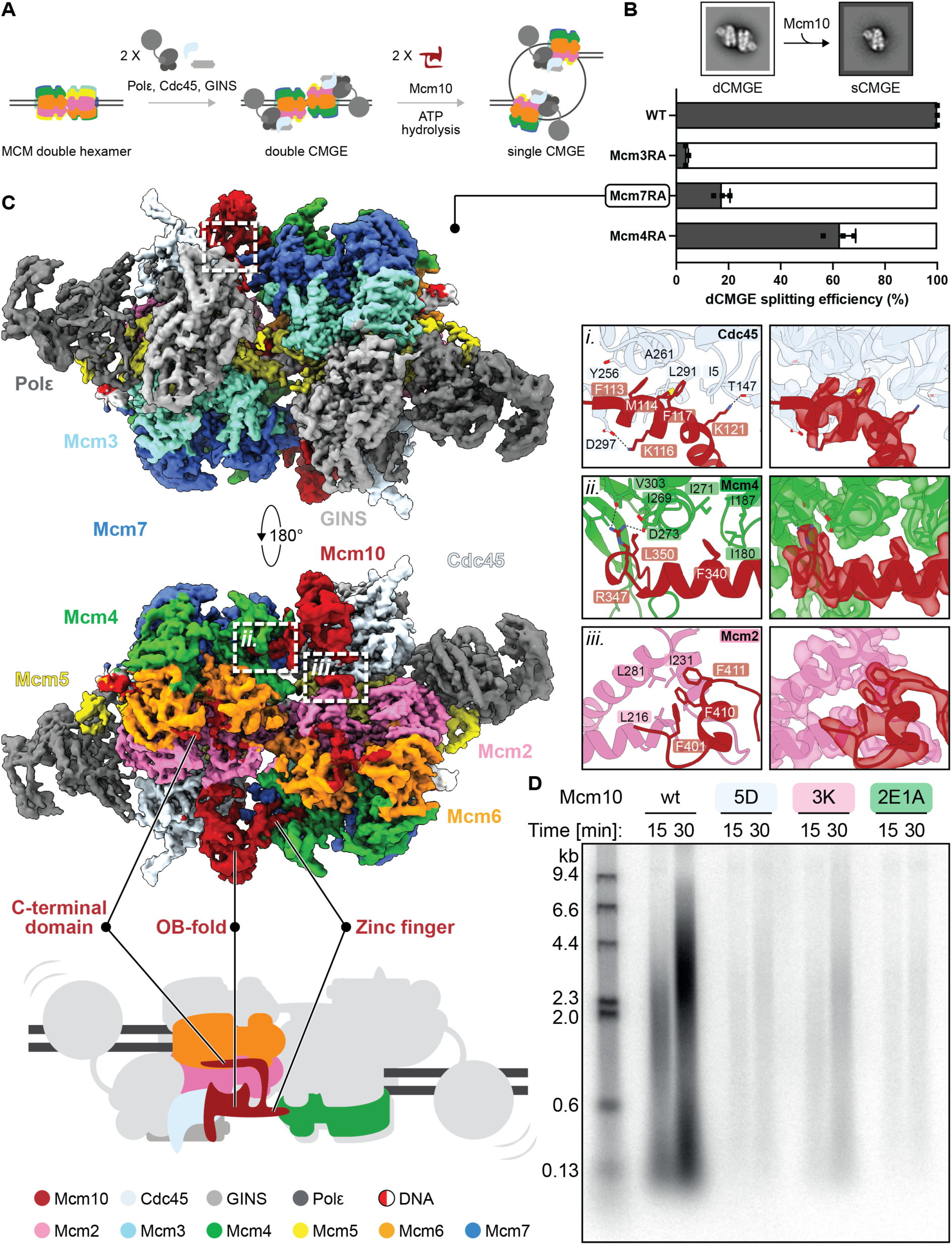
Mcm7RA traps double CMGE-Mcm10. (**A**) Cartoon depicting the sequence of events leading to CMGE activation. First, a DNA-loaded DH is converted to dCMGE upon recruitment of Cdc45, GINS and DNA polymerase epsilon. Then, Mcm10 triggers ATP hydrolysis by CMGE, leading to the ejection of the lagging strand template and the establishment of diverging replication forks. (**B**) Bar graph comparing dCMGE splitting efficiency by Mcm10 using Mcm2-7 wild type (wt), Mcm3RA, Mcm7RA or Mcm4RA. This experiment was performed three times. (**C**) Cryo-EM map of dCMGE10^Mcm7RA^. The N-terminal OB domains from two Mcm10 subunits capture converging CMGEs in a defined register, by bridging across Cdc45 and Mcm4 from opposing CMGE monomers. (**C-i**) The first tethering element of Mcm10 on CMG is an alpha helix (residues 113-121) that binds to Cdc45. (**C-ii**) The second tethering element is an alpha helix insertion of the OB fold (residues 333-350) contacting Mcm4. (**C-iii**) A prominent contact element between the C-terminal domain of Mcm10 and CMG is formed by three phenylalanines (F401, F410, F411) that insert into a hydrophobic pocket on Mcm2. (**D**) Structure-guided Mcm10 mutants block or severely impair DNA replication in reconstituted reactions with purified proteins.

Strikingly, reactions containing Mcm7RA and Mcm10 yielded a distinct class of dCMGE averages in which the N-terminal MCM tiers were more tightly interacting than in any previously observed configuration (*15*) (fig. S1A). We reasoned that this unusual architecture represents a stalled intermediate on the pathway to Mcm10-dependent helicase activation, and decided to image the purified particles by cryo-EM.

### Cryo-EM structure of dCMGE10^Mcm7RA^

We solved a structure of dCMGE assembled using Mcm7RA and Mcm10 (dCMGE10^Mcm7RA,^ fig. S2, 3). The N-terminal OB domains from two separate Mcm10 subunits tether the opposing CMGE monomers, each bridging Cdc45 and Mcm4 across a tightened interface of the helicase dimer. The C-terminal Mcm10 domain decorates the outer perimeter of Cdc45, Mcm2 and 6, nestling into a groove formed between the N-terminal and ATPase tiers of each MCM hexamer (Fig. 1C).

A structural role for the essential N-terminal OB fold of Mcm10 in reshaping the dCMGE interface was unexpected. Compared to ATP-dCMGE (Fig. 2A) assembled in the absence of Mcm10 with wild type proteins (*15*), the two CMGEs appear to have converged. Two tethering elements are poised to stabilise the tightened dimer. The first, conserved across fungi, is a Cdc45-contacting element featuring five Mcm10 residues, engaged in a set of hydrophobic and polar interactions (Fig. 1C-i). The second tethering element is an Mcm10 alpha helix insertion of the OB fold contacting Mcm4. Here, three universally conserved Mcm10 residues access a groove in Mcm4 of the opposed CMGE. This MCM epitope is known to become exposed when DH is phosphorylated by DDK (*9, 10*), on the path to dCMGE formation (*24*) (Fig. 1C-*ii*). Both interactions are essential for function. In fact, Mcm10^5D^ (F113D, M114D, K116D, F117D, K121D) and Mcm10^2E1A^ (R347E, F340E, L350A), which target the Cdc45 and Mcm4 contacts respectively, block stable dCMGE10 assembly with Mcm7RA, severely impair Mcm10-dependent dCMGE splitting (fig. S1B-F), and abolish DNA replication initiation in reconstituted reactions (Fig. 1D).

**Fig. 2.**
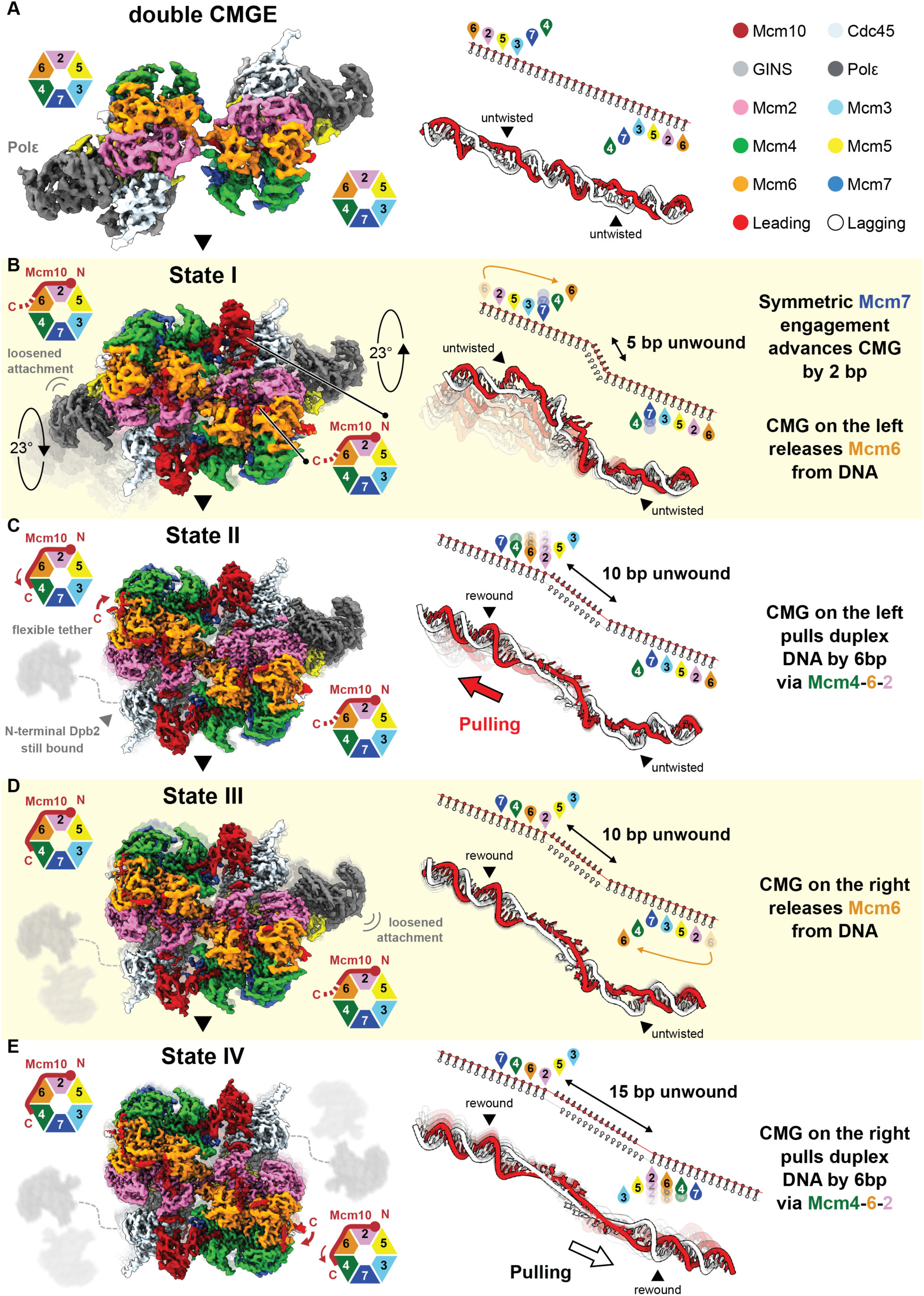
Four intermediates of Mcm10-mediated origin DNA shearing. (**A**) Cryo-EM map of dCMGE (PDB 7Z13(*15*)) and cartoon of the two MCM ATPase hexamers. The DNA model is shown below a cartoon depicting the engagement state of MCM ATPase pore loops. Both CMGEs engage DNA via Mcm6, 2, 5, and 3 and the duplex is untwisted within the ATPase tiers of each CMGE. (**B**) Cryo-EM map of dCMGE10^Mcm7RA^ State I. Mcm10 binding induces the CMGs to rotate 23° in a direction opposite the DNA twist, resulting in 5 bp unwound at the dimerisation interface. Each CMG has advanced by 2 bp via Mcm7, and CMG on the left has released Mcm6 from DNA, resulting in loosened attachment of Pol epsilon from CMG. Mcm10 C-terminal domain seals the Mcm6-Mcm2 ATPase interface. (**C**) Cryo-EM map of dCMGE10^Mcm7RA^ State II. CMG on the left has pulled duplex DNA by 6 bp via Mcm4, 6, and 2, resulting in rewinding of the untwisted DNA within its ATPase tier and further DNA opening at the dimer interface (10 bp unwound). Pol epsilon is flexibly tethered to CMG on the left, with N-terminal Dpb2 as the only element still visible, while C-terminal Mcm10 staples Mcm6 and Mcm4 together. (**D**) Cryo-EM map of dCMGE10^Mcm7RA^ State III. CMG on the right has released Mcm6 from DNA, resulting in loosened attachment of Pol epsilon from CMG. (**E**) Cryo-EM map of dCMGE10^Mcm7RA^ State IV. CMG on the right has pulled duplex DNA by 6 bp via Mcm4, 6, and 2, re-establishing the symmetry of the complex and further unwinding DNA at the dimer interface (15 bp unwound).

Mcm10 is known to stimulate ATPase driven DNA unwinding and fork rates in replication assays (*21, 25*). We previously demonstrated that the fork rate enhancement function is contained in the C-terminal Mcm10 domain. While we could not explain the mechanism at the time, we showed that this function is dispensable for replication initiation (*16*). Our current observation that C-terminal Mcm10 straddles the Mcm2 and Mcm6 subunits suggests how replisome rate stimulation could be achieved. In fact, bringing AAA+ subunits in close proximity is an established mechanism to enable the reconstitution of a catalytically active ATPase site (*26*), which would promote fork advancement. To test this model, we set out to disrupt the Mcm10-Mcm2-Mcm6 tripartite interaction. One prominent tethering element is formed by three phenylalanines in C-terminal Mcm10 (F401, F410, F411, Fig. 1C-iii), which insert into a hydrophobic pocket on the Mcm2 surface. Breaking this contact with three lysine substitutions, engineered in the Mcm10(3K) variant, yielded a striking phenotype, though milder compared to the dead N-terminal Mcm10 variants described above. While stable dCMGE10^Mcm7RA^ assembly was still blocked, in fact, only a partial dCMGE splitting defect could be observed, and some replication signal was still detected in the *in vitro* reconstituted assay, with a shorter nascent DNA product consistent with defective fork rate (Fig. 1D and fig. S1B-F). Together, our data agree with our previous finding that fork rate enhancement by C-terminal Mcm10 is dispensable for replication initiation (*16, 27*).

So far, we have described the structure of dCMGE10^Mcm7RA^ where Mcm10 locks two converging dCMGE helicases in a defined register. N-terminal Mcm10 contacts established with this Mcm7 variant support replication initiation with wild type proteins, indicating that this structure reflects a *bona fide* intermediate of the origin activation reaction. Also, C-terminal Mcm10 sealing together two MCM motor subunits further provides a mechanism for the (dispensable) fork-rate stimulation function of Mcm10.

### Four ATPase intermediates in dCMGE10^Mcm7RA^

We analysed the new dCMGE10^Mcm7RA^ structure with three-dimensional classification, seeking to understand how Mcm10 can affect catalysis in the two converging CMGEs. We observed different ATPase states, reflected in gross reconfigurations of Pol epsilon and Mcm10 tethering to the two CMGs, as well as different DNA grips inside the MCM pore (fig. S2, 3). In State I (4.2 Å resolution) Dpb2 and C-terminal Pol 2 of Pol epsilon are fully engaged to one CMGE (on the right in Fig. 2B), similar to dCMGE (*15*) (Fig. 2A). Pol epsilon on the opposed CMG, instead, is partially disengaged with C-terminal Pol 2 distanced from Cdc45 (on the left in Fig. 2B). In State II (3.4 Å resolution), the right Pol epsilon is still fully engaged to CMG, whereas the polymerase on the left is almost completely detached, with only the small helical N-terminal Dpb2 domain visible tethered to GINS (Fig. 2C). In State III (4 Å resolution), the density for the right polymerase becomes partially degraded, reflecting loosened attachment (Fig. 2D). State IV (3.5 Å) is symmetric and N-terminal Dpb2 is the only Pol epsilon element visible on either of the two CMGEs (Fig. 2E).

A second conformational change becomes evident as Pol epsilon disengages from tight CMG binding (Fig. 2C-E): a C-terminal Mcm10 alpha helix (residues 534-552) becomes visible, stapling Mcm6 and Mcm4 together. This observation reinforces the idea that straddling ATPase subunits underlies the fork-rate stimulation function of C-terminal Mcm10 (*21, 25*). We also note that, with half of the MCM ring now sealed by this C-terminal Mcm10 element (Fig. 2E), neither the Mcm6-2 nor the Mcm4-6 intersubunit interfaces are likely to open. This excludes the possibility that either of these two interfaces serves as the lagging strand ejection gate, at least for as long as they remain Mcm10-associated.

The degree of Pol epsilon engagement by CMG does not only come with changes in Mcm10 binding. Also different DNA gripping modes inside the MCM pore can be observed (*14, 28*) (Fig. 2 and Fig. 3). When Pol epsilon is fully engaged to CMG, in fact, conserved lysines in the Pre-sensor 1 (PS1) pore loops of Mcm6, 2, 5, and 3 track along duplex DNA by binding the phosphodiester backbone of the leading-strand template (as observed in dCMGE, Fig. 2A and 3A). When Pol epsilon becomes loosely attached, Mcm7 engages the leading strand template at the front of the helicase, followed by Mcm6 disengaging DNA at the back (as observed in States I, II and III, Fig. 2B-D and Fig. 3B-D). When Pol epsilon is only tethered via N-terminal Dpb2, PS1 pore loops of Mcm7, 4, 6, and 2 are leading-strand-bound (observed in states II-IV, Fig. 2C-E and 3C-E).

**Fig. 3.**
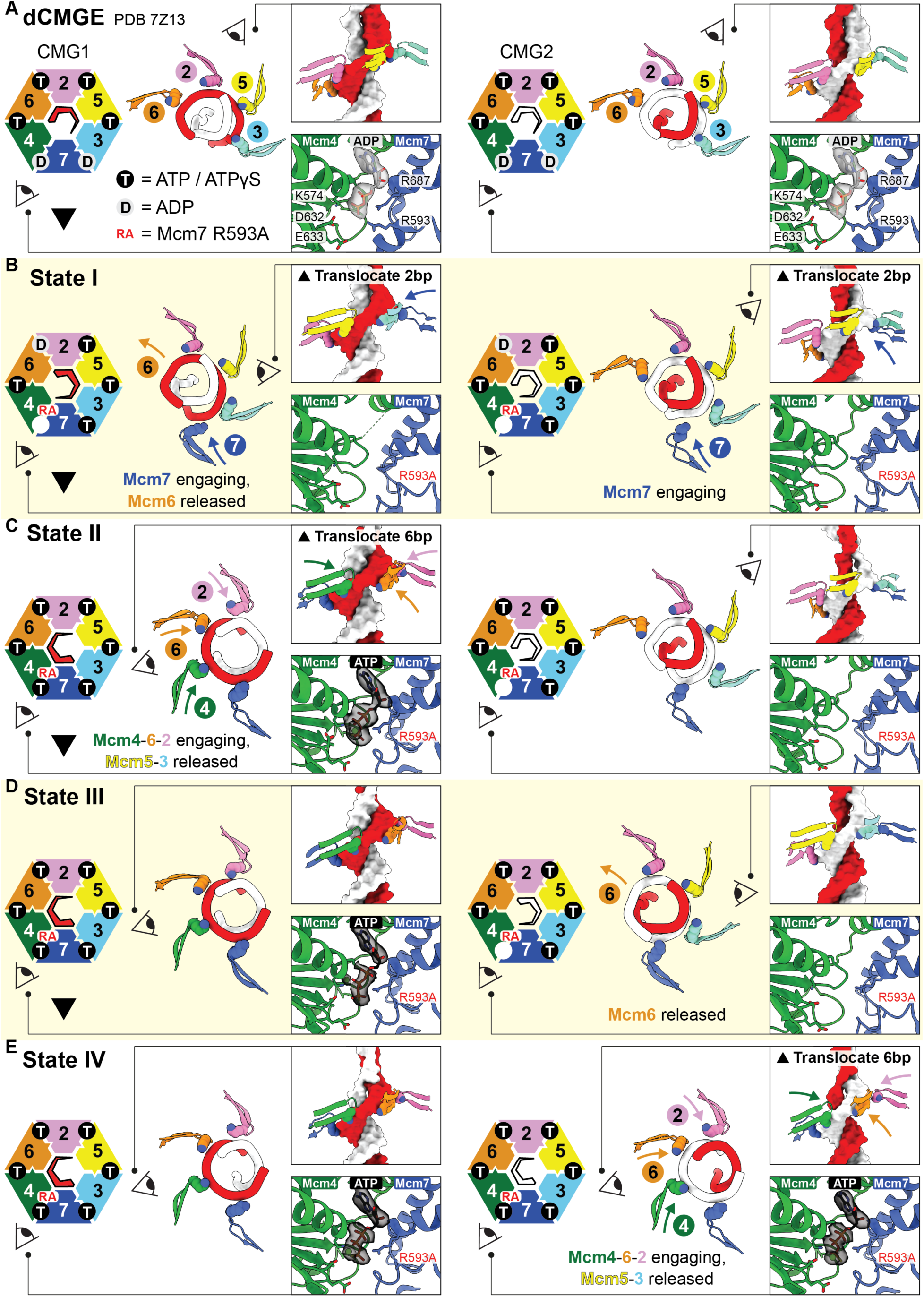
The mechanism of origin melting by the MCM ATPase. (**A**) DNA binding in the dCMGE. Top view cartoon of the MCM ATPase hexamer next to the atomic model of the pore loops bound to DNA. The PS1 lysines that contact the duplex DNA backbone are shown as spheres. The two insets highlight the pore loop staircase and the Mcm4/Mcm7 ATP-binding interface with nucleotide density shown as transparent surface. Both CMGEs engage DNA via Mcm6, 2, 5, and 3. (**B**) DNA binding in dCMGE10^Mcm7RA^ State I. Both CMGs translocate by 2 bp via Mcm7 engagement. CMG on the left also releases Mcm6 from DNA. Further DNA translocation via Mcm4 is impaired by the Mcm7RA mutation that prevents nucleotide binding at the Mcm4/Mcm7 interface. (**C**) DNA binding in dCMGE10^Mcm7RA^ State II. CMG on the left pulls DNA by 6 bp as it engages the duplex via Mcm4, 6, and 2 and releases the Mcm5 and 3 contacts. Further DNA pulling is impaired by the Mcm7RA mutation that prevents ATP hydrolysis at the Mcm4/Mcm7 interface, thereby preventing disengagement of Mcm7 from DNA. (**D**) DNA binding in dCMGE10^Mcm7RA^ State III. CMG on the right releases Mcm6 from DNA. (**E**) DNA binding in dCMGE10^Mcm7RA^ State IV. CMG on the right pulls DNA by 6 bp as it engages duplex DNA via Mcm4, 6, and 2 and releases Mcm5 and 3. Further DNA pulling is impaired by the Mcm7RA mutation that prevents ATP hydrolysis at the Mcm4/Mcm7 interface, thereby preventing release of Mcm7 from DNA.

Previous studies on helicase translocation led to the proposal of a hand-over-hand model, where ATP binding by neighbouring subunits promotes sequential pore-loop-DNA engagement at the front of the helicase, while hydrolysis drives disengagement at the back (*29–32*) (Movie S1). Key elements of this rotary mechanism agree with our dCMGE observations (*15*) and the four dCMGE10^Mcm7RA^ states described in this study, where the inactivating change in the Mcm7 Arginine Finger is expected to be partially defective in either binding or hydrolysis of ATP. Here is why. In ATP-dCMGE, Mcm6, 2, 5, and 3 PS1 pore loops engage DNA (Fig. 2A and Fig. 3A). Upon ATPase activation by Mcm10, an ATP-binding defect at the Mcm4-7^RA^ ATPase site would block MCM motor advancement with the Mcm7 PS1 pore loop bound to DNA at the front of the helicase. Consistently, in States I, II and III where Mcm7 binds DNA in front, no nucleotide is observed in the Mcm4-7^RA^ ATPase site (both CMGEs in Fig. 2B and Fig. 3B, right-hand CMGE in Fig. 2C-D and Fig. 3C-D). Instead, in States II, III and IV, defective hydrolysis in the Mcm4-7^RA^ ATPase site prevents Mcm7 from releasing DNA at the back of the helicase, stabilising a Mcm7-4-6-2 DNA grip (left-hand CMGE in Fig. 2C-D and Fig. 3C-D and both CMGEs in Fig. 2E and Fig. 3E, and Movie S1). As predicted by the hand-over-hand model (*29–32*), ATP density (possibly ATPγS, which was used for the final elution step in our preparation) can be observed sandwiched between Mcm4 and Mcm7^RA^ in these States (Fig. 3C-E).

### ATP-hydrolysis driven DNA shearing by converging CMGEs

We next describe how ATPase controlled DNA translocation in the Mcm10-constrained dCMGE^Mcm7RA^ causes DNA opening at the MCM homodimerisation interface. Our previous work showed that the ATP-bound dCMGE assembly promotes local untwisting of duplex DNA inside the two MCM ATPase tiers, while the DNA segment trapped in between the partially-separated helicases retains a B-form structure (Fig. 2A) (*15*). Mcm10 engagement, as observed in dCMGE10^Mcm7RA^ State I, induces two structural transitions. First, the two MCM motors converge, with each helicase advancing by two base pairs, through DNA engagement by Mcm7 ahead of the helicase. This generates torsional strain on the double helix in front (Fig. 2B and Fig. 3B). Second, by locking the two CMGEs together, the N-terminal domain of Mcm10 entraps a stretch of underwound DNA focused between the two converging helicases (Fig. 2B). Here, Mcm10 binding across the two CMGEs induces a rotation in the CMGs in a direction opposite the duplex DNA twist, resulting in underwinding of the double helix at the dimer interface (Fig. 2B and Fig. 4A). As they transition from dCMGE to dCMGE10^Mcm7RA^ State I, the two Zinc Fingers of Mcm2 across the CMGE dimer come in close proximity, generating a steric barrier to B-form DNA stability, which forces the double helix to flatten and open (Movie S2). The torsional strain stored in this structure is resolved through the disruption of five base pairs (Fig. 2B), with three bases per strand flipped out symmetrically around the centre of the CMGE dimer (Fig. 2B and Fig. 4B).

**Fig. 4.**
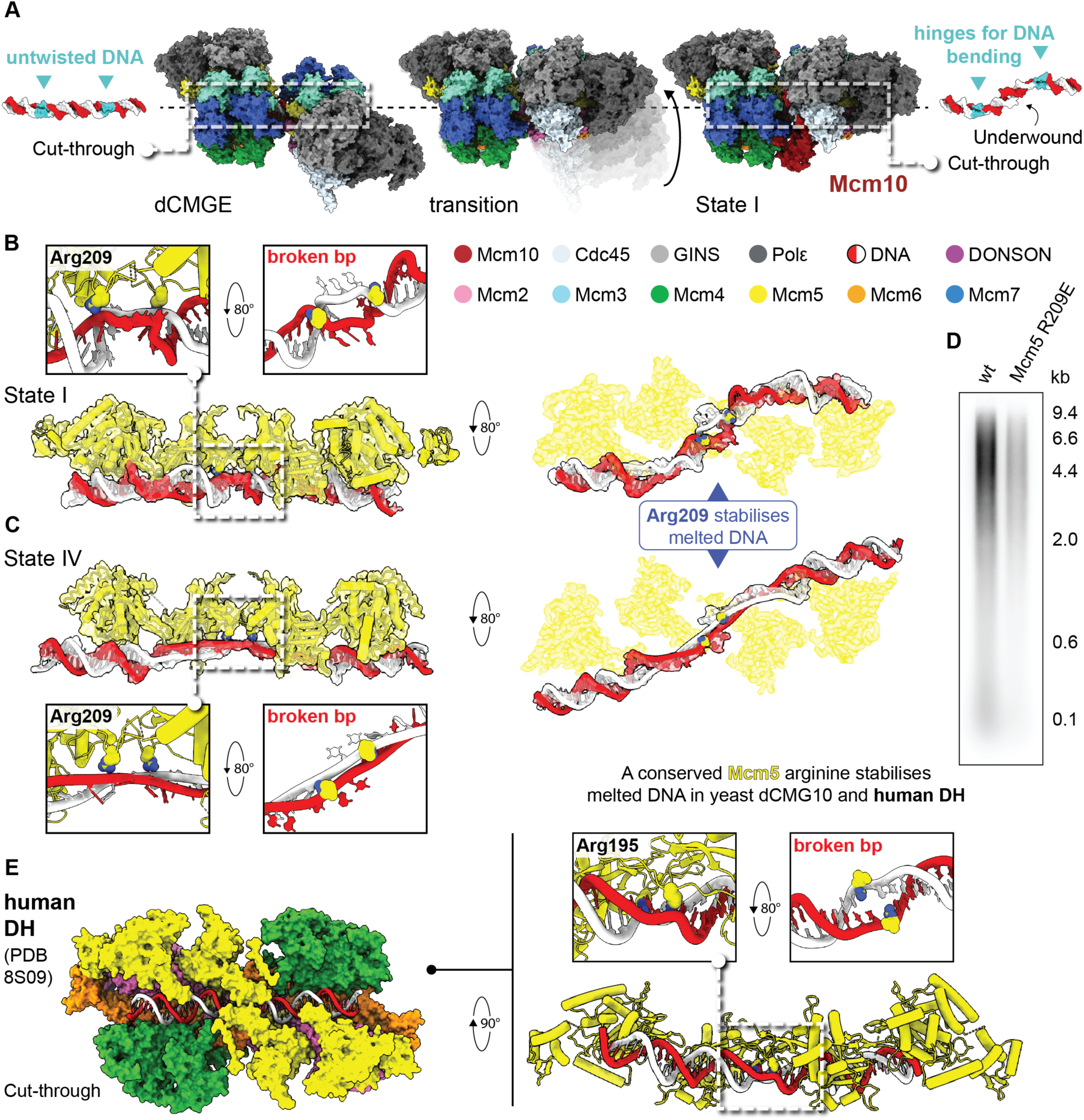
The melted origin is stabilized by a conserved Mcm5 element. (**A**) Mcm10 binding across the two CMGEs promotes a rotation opposite the DNA twist, resulting in underwinding of the double helix. The regions of DNA that were untwisted in the dCMGE become the hinges that accommodate the bending of the DNA molecule upon transition to dCMGE10^Mcm7RA^ State I. (**B**) Melted DNA in dCMGE10^Mcm7RA^ State I is stabilised by Mcm5 Arg209 engaging the phosphodiester backbone on either side of the open bubble. Mcm5 Arg209 is shown as spheres in the model. (**C**) The same Mcm5 Arg209-DNA contact persists through State IV where DNA untwisting is maximised. (**D**) Charge-reversal of Mcm5 Arg209 to glutamate causes a severe DNA replication initiation defect in reconstituted *in vitro* reactions. (**E**) In the human system, DNA untwisting is achieved upon DH loading. During this process breakage of one base pair is stabilised by one universally conserved residue, Mcm5 Arg195, which corresponds to Arg209 in yeast Mcm5.

Upon transition from State I to State II, one of two CMGEs moves forward by three subunits (6 bp, Fig. 2C and Fig. 3C). As DNA translocation occurs, the DNA that was untwisted within one ATPase tier (Fig. 2A and Fig. 2B) becomes rewound (CMGE on the left, Fig. 2C). This creates positive twist behind the helicase, which is compensated by negative twist in front, promoting further DNA opening between the two converging helicases (Fig. 2C). Transition from State II to State III involves a minor rearrangement, with Mcm6 disengaging from DNA at the back of the helicase and no change in the structure of DNA (Fig. 2C and Fig. 2D). Transition from State III to State IV (Fig. 2D and Fig. 3D) involves a three-subunit translocation from the second CMGE such that symmetry is restored in the dimeric assembly (Fig. 2E and Fig. 3E). By the time State IV is established, a 15-bp DNA segment has become untwisted (Fig. 2E and Fig. 4C). Base pairing in this entire DNA segment appears out-of-register, as would happen when leading strand translocation from one CMGE extracts the lagging strand from the opposed CMGE in the dCMGE10^Mcm7RA^ structure and vice versa (Fig. 2E, Fig. 4C and Movie S3). This is similar to the DNA shearing mechanism previously envisaged for the SV40 Large T antigen (*33*) and the CMG helicase (*34*).

New contacts are created in the Mcm10-braced CMGE dimer, which stabilise the sheared DNA. For example, Mcm5 Arg209 stabilises the flipped-out bases by engaging the adjacent phosphodiester backbones on either side of the melted DNA bubble. Established in State I, this configuration persists through State IV where DNA untwisting is maximised (Fig. 4B and Fig. 4C). An MCM variant bearing a single reverse charge mutation to glutamate, Mcm5(R209E), can load DHs on DNA to wild type levels (fig. S1G), but yields a severe initiation defect, observed in the reconstituted DNA replication assay (Fig. 4D). This result indicates that the unwound DNA structure observed in dCMGE10^Mcm7RA^ reflects a *bona fide* initiation intermediate and not a dead-end state due to the engineered ATPase defect (Mcm7RA).

Unlike in yeast, human DH loading is sufficient to untwist the double helix and break one base pair in the DNA entrapped between the two MCM hexamers (*35, 36*). We note that the Mcm5 ZnF Arginines that stabilise broken DNA base pairing in the human DH (Arg195, Fig. 4E) and the yeast dCMGE10^Mcm7RA^ (Arg209, Fig. 4B and Fig. 4C) are universally conserved (*36*). Thus, while the order of events that achieve strand separation at replication origins has diverged during evolution, the residues that engage melted DNA have been retained from yeast to human.

Because this work relies on ATPase mutants, altered translocation kinetics should be considered when interpreting our structures. The asymmetric states we observe, where one CMGE translocates before of the other, may reflect an inherent feature of the melting mechanism or a consequence of inefficient translocation due to the use of a defective ATPase variant. Our current data do not distinguish between these possibilities.

### Lagging strand extracted through Mcm2-5

While duplex DNA is known to be threaded through the Mcm2-5 opening during MCM loading (*37–40*), it remains unknown whether lagging strand ejection occurs through this same DNA gate or whether multiple MCM interfaces can serve as alternative exit sites. Inspection of the various dCMGE conformers reveals how Mcm10 engagement, and the ensuing DNA translocation, commit the lagging strand to ejection through one specific gate. In fact, transition from ATP-dCMGE (*15*) to dCMGE10^Mcm7RA^ State I reveals that the lagging strand becomes symmetrically entrapped within the ZnF domain of Mcm2 and Mcm5 of each CMGE helicase (Fig. 5A-C). As described above, the transition to downstream states State II to IV involves duplex DNA threading from N- to C-terminal MCM, through the ATPase hexamer (Fig. 2C-E and Fig. 3C-E). To feed enough DNA to sustain translocation by each MCM motor, the lagging strand is extracted from the opposed hexamer and further threaded through a gap between the ZnF domains of Mcm2 and Mcm5 (Fig. 5D and Fig. 5E and Movie S4). Unidirectional DNA translocation by helicase motors is known to depend on ATP hydrolysis, which is an irreversible chemical process (*41*) (Fig. 5D). Hence, DNA committed to and extracted through a gap within the 2-5 gate can become fully ejected only through the complete opening of the same 2-5 interface. In fact, for a different gate to be operated the helicase would need to run in reverse, which is something the MCM motor does not do.

**Fig. 5.**
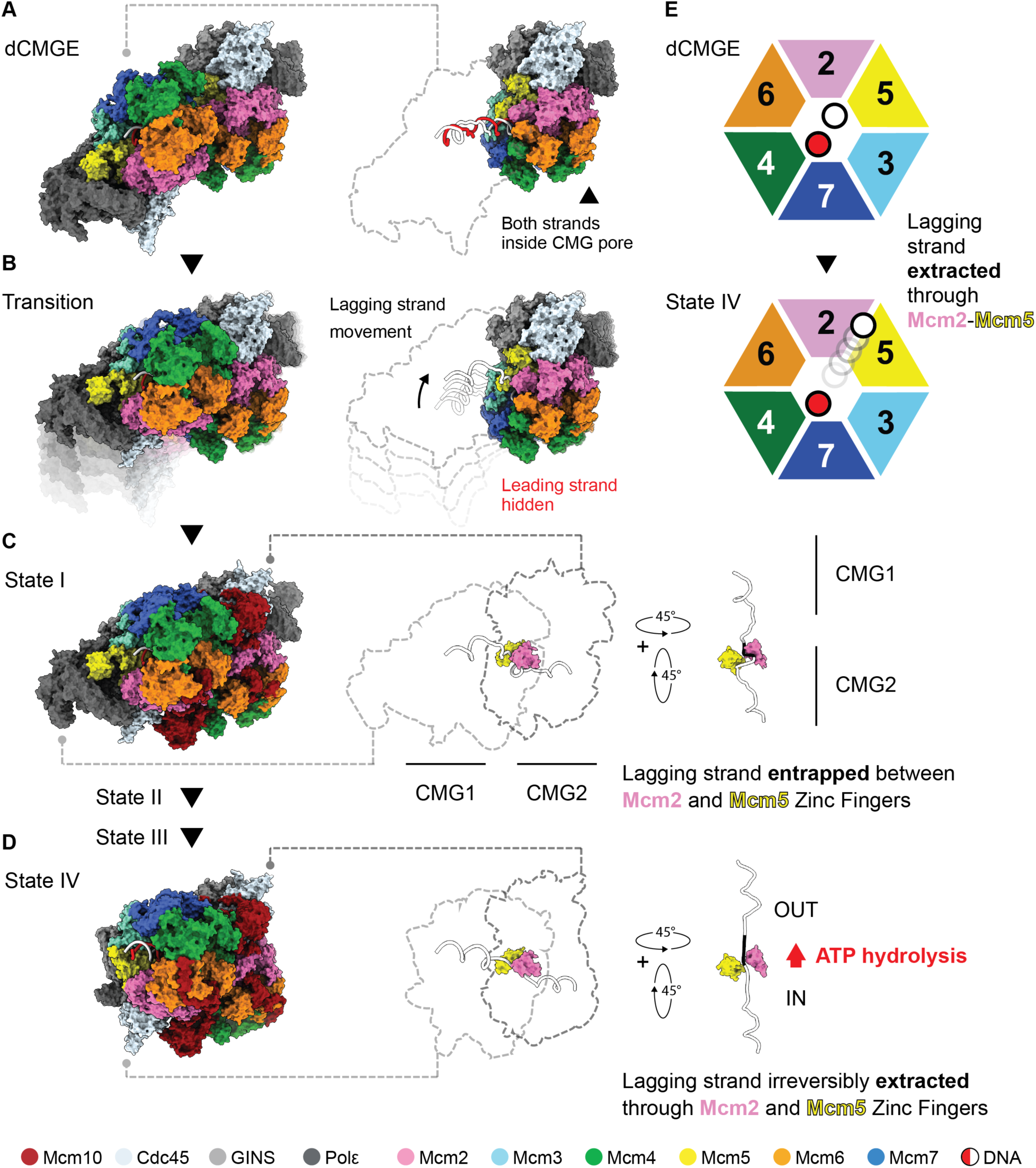
The lagging strand template is extracted through the Mcm2-5 gate. (**A**) Cryo-EM structure of the dCMGE, where both DNA strands are inside the CMG pore. (**B**) Upon transition to dCMGE10^Mcm7RA^ State I, the lagging strand template moves towards the interface between the ZnF of Mcm2 and Mcm5. (**C**) In dCMGE10^Mcm7RA^ State I, the lagging strand is entrapped between the ZnF of Mcm2 and Mcm5. (**D**) In dCMGE10^Mcm7RA^ State IV, ATP hydrolysis driven translocation has irreversibly extracted the lagging strand template through the ZnF of Mcm2 and Mcm5. (**E**) Top-view cartoon representation of the MCM ATPase in the dCMGE and in State IV, depicting the commitment of the lagging strand template to ejection from the Mcm2-5 gate, upon Mcm10-triggered ATP hydrolysis.

A second, independent observation points to opening of the Mcm2-5 gate as the sole possible step following State IV. By this stage, the lagging strand is fully stretched between the two MCM hexamers (Fig. 5D), meaning the opposed helicase has no further DNA to supply. With DNA fully extended, one additional translocation step (Mcm4 engaging DNA ahead of Mcm7) would presumably destabilise the dCMGE10^Mcm7RA^ structure. We reasoned that, to progress beyond State IV, the Mcm2-5 gate must open, so that the lagging strand can be ejected, allowing each MCM motor to continue translocating past each other as an uncoupled sCMGE.

To test this scenario, we imaged the dCMGE splitting reaction using an ATPase-inactivating variant (Mcm4RA), targeting the next active site to be fired according to the hand-over-hand translocation model.

### Cryo-EM analysis of sCMGE10^Mcm4RA^

We assembled the origin activation reaction with Mcm10 and Mcm4 RA, bearing an inactivating mutation in the ATPase subunit proximal to Mcm7. We established earlier that Mcm4 RA robustly supports dCMGE splitting into two single CMGEs (Fig. 1B and fig. S1A). Solving the cryo-EM structure revealed that Mcm10 disengages from the CMGE when two helicases become separated (Fig. 6A and fig. S5). Two different sCMGE species can be observed, one encircling duplex and the other single-stranded DNA (fig. S5F). We reasoned that the duplex-interacting species might reflect failed lagging strand ejection due to the defective ATPase mutant used in our reaction. Because our aim was to gain further insight into the lagging strand ejection process, we focused our attention on the subset of sCMGE complexes that successfully ejected the lagging strand and only engage single-stranded DNA within the MCM channel. On the N-terminal side of the MCM motor of sCMGE^Mcm4RA^, we observed duplex DNA capped by covalently linked MH. This stretch of double helix becomes unwound as it enters the MCM motor (Fig. 6A and fig. S4H), while the leading-strand template traverses the ATPase central channel and contacts PS1 pore loops of Mcm6, 2, 5, and 3. This is the same leading-strand engagement reported for sCMGE assembled with Mcm10 and wild type MCM (*16*). Duplex DNA becomes rewound as it emerges from the MCM ATPase pore, while the lagging strand can be observed sitting on the outer MCM surface. In this way, a stretch of double helix at both the front and the back of the helicase are connected by continuous leading and lagging strand density, forming a complete DNA bubble topologically entrapped by the MCM ring in sCMGE10*^Mcm4RA^* (Fig. 6A). The lagging strand ejected from MCM rests within a positively charged groove spanning the N- and C-terminal domains of Mcm2 on one side and Cdc45 on the other (Fig. 6B). This lagging-strand engagement site had not been observed before and is distal from the Mcm3-5 groove through which the lagging strand is channelled in the fully established replication fork (*29, 42*). Two observations indicate that this structure likely reflects a DNA engagement intermediate immediately following strand ejection. First, the lagging strand binding site in sCMGE^Mcm4RA^ is closest to the Mcm2-5 gate from which it was ejected (Fig. 6A). Second, when superposing dCMGE10^Mcm7RA^ to sCMGE^Mcm4RA^, the single-stranded binding OB fold of Mcm10 is poised right next to the Mcm2-5 gate and is located in the immediate vicinity of the ejected DNA, bound by Mcm2 in our sCMGE10*^Mcm4RA^* structure. The binding polarity of the Mcm10 OB fold had been previously established with a frog Mcm10-DNA co-crystal structure from Brandt Eichman’s lab (*20*) and agrees with our structure of the open bubble (Fig. 6C and Fig. 6D). We note that this directionality supports selective lagging strand engagement by Mcm10, upon DNA ejection (Fig. 6E).

**Fig. 6.**
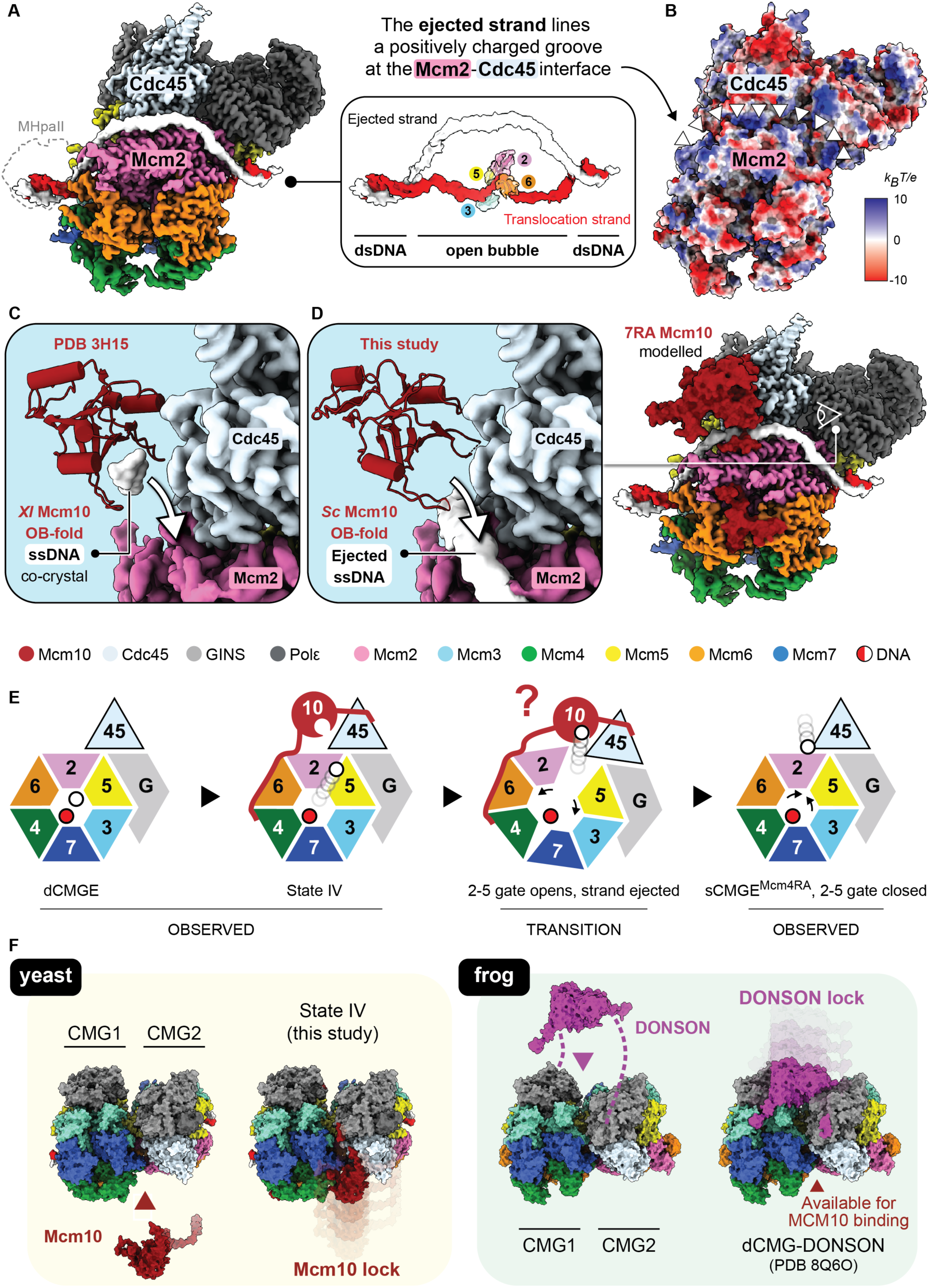
Post-ejection sCMGE captured with Mcm4RA. (**A**) Cryo-EM map of sCMGE10^Mcm4RA^. Duplex DNA at the N-terminal side of MCM becomes unwound as the leading strand template enters the MCM pore, while the lagging strand lines the outer Mcm2-Cdc45 interface. Duplex DNA rewinds at the C-terminal side of MCM where the leading strand template emerges from the MCM ATPase pore. (**B**) The ejected strand rests within a positively charged groove at the interface between Mcm2 and Cdc45, closest to the Mcm2-5 gate. (**C**) The crystal structure of *Xenopus laevis* Mcm10 OB-fold domain bound to ssDNA (PDB 3H15(*20*)) was superposed to our dCMGE10^Mcm7RA^ structure. The binding polarity of the Mcm10 OB-fold supports selective engagement of the ejected strand. (**D**) The structure of dCMGE10^Mcm7RA^ was superposed to sCMGE10^Mcm4RA^. The ssDNA-binding groove of Mcm10 is poised to capture the strand ejected from the Mcm2-5 gate and deliver it to the Mcm2-Cdc45 interface upon Mcm10 eviction. (**E**) Top-view cartoons of the CMG depicting the model for Mcm10-dependent lagging strand template ejection. In the dCMGE, both DNA strands are inside the CMG pore. Upon Mcm10 engagement and activation of the ATP hydrolysis function, the lagging strand is irreversibly extracted through the Mcm2-5 interface, such that gate opening would complete ejection. The Mcm10 OB-fold is poised to capture the ejected strand and deliver it to the Mcm2-Cdc45 interface upon Mcm10 eviction. (**F**) The way two converging yeast CMGs are locked by Mcm10 recapitulates the architecture observed for the frog dCMG locked bound by dimeric DONSON. While yeast and frog CMGs can be perfectly superposed, Mcm10 and DONSON bind on opposite sides of the complex. Thus, in the frog dCMG, DONSON and MCM10 engagement might occur concomitantly.

We also note a technical correction. A density on the N-terminal face of the helicase in our sCMGE structure, which we conclusively identified as covalently linked MH, was tentatively attributed to Mcm10 in a previously published, lower resolution structure (*16*).

## Discussion

Accumulating evidence indicates that two key events during origin activation are mechanistically coupled: the physical separation of two fully formed replicative helicases (the dCMGE-to-sCMGE transition), and the symmetric ejection of the two lagging strand templates that establishes divergent replication forks. A first indication of this coupling comes from work by Steve Bell’s group, which showed that monomeric CMGs can be assembled at an origin of replication and can untwist duplex DNA, yet fail to eject the lagging strand during the final Mcm10-dependent step of helicase activation (*43*). A second indication comes from our recent discovery that dCMGEs can form in the absence of Sld2, but their splitting into sCMGEs is defective. Notably, the minority of sCMGEs that do form under these conditions fail to eject the lagging strand despite the presence of Mcm10 (*14*). Our current study explains the underlying mechanism. We employed an ATPase-defective MCM variant (Mcm7RA) to slow the origin DNA melting reaction, enabling us to capture four elusive structural intermediates. We find that after partially separating within the dCMGE structure, two CMGEs translocate toward one another and become interlocked, with two Mcm10 molecules restraining their converging N-terminal MCM domains in a fixed position. Because Mcm10 activates ATPase-powered DNA translocation (*25*), the two helicases continue threading duplex DNA from the N- to the C-terminal side of MCM, thereby shearing the duplex DNA trapped between the two helicase motors. While each CMGE primarily tracks on the leading strand template, the lagging strand is passively extracted from each MCM by the ATPase-powered pulling action of its cognate CMGE. During this process, the Mcm2–5 inter-subunit interface is selected as the passage through which the lagging strand is extracted. At this stage the DNA is fully stretched, such that additional translocation would drive separation of the two helicases, at which point they would both complete lagging strand ejection, split into individual sCMGEs, release Mcm10, and cross paths. To visualise the post-separation state, we employed a second ATPase-defective variant (Mcm4RA), which allowed us to capture a CMGE assembly that has fully separated from its cognate helicase and transitioned to encircling the leading strand template alone. In this structural intermediate, the lagging strand is observed resting just outside the Mcm2–5 interface, corroborating the conclusion that this gate is used for strand ejection and replication fork establishment. Together, our observations explain that MCMs are loaded at origins as symmetric double hexamers because they extract DNA from one another once they become activated and this establishes two divergent replication forks (*3, 4*). Our results also reconcile the established hand-over-hand mechanism for DNA translocation that supports replication fork unwinding (*29, 32*) with the mechanism of duplex DNA melting.

Finally, our structures of two convergent yeast CMGEs locked by Mcm10 provide unexpected insights into the mechanism of origin activation in higher eukaryotes. Replicative helicase maturation in metazoans requires DONSON, a replisome assembly factor lost during yeast evolution (*44–48*). Like Dpb11 in yeast (*14*), DONSON recruits GINS to MCM to promote CMG assembly (*44–48*), while also reshaping the N-terminal dimerisation interface of the DH to generate a CMG dimer bridged by dimeric DONSON (*47*). The structure of a CMG dimer observed in dCMGE10^Mcm7RA^ is strikingly similar to that of the DONSON-stabilised CMG dimer (RMSD 4.5 Å) (Fig. 6F). This observation suggests that in metazoans duplex DNA at replication origins might become sheared by two converging CMGEs like in the yeast system. DONSON might function to lock the two CMGEs like Mcm10. Consistent with this, concomitant binding of DONSON and Mcm10 to a converged CMG dimer would be physically allowed, suggesting these two firing factors might act in the same step during origin firing, and potentially play a partially redundant function. In this respect, while Mcm10 is essential for replication initiation *in vitro* and for cell viability in yeast, work in *C. elegans* has established that DONSON, but not Mcm10, is required for viability (*44*), suggesting that DONSON serves a prominent role in duplex DNA melting during metazoan origin activation.

## Materials and Methods

### Protein Expression and Purification

The following proteins were expressed and purified according to established protocols: MH, ORC, Cdc6, Mcm2-7/Cdt1 (wild-type, 3RA, 7RA, 4RA), DDK, CDK, Sld3/7, Cdc45, GINS, Polε, Mcm10, Replication Protein A (RPA), Topoisomerase I (TopoI), Polα (*15, 17, 49–53*). Protein variants bearing point mutations were prepared using the wild-type purification protocol unless otherwise noted.

### Cloning, expression and purification of Mcm2-7/Cdt1 (Mcm5-R209E)

Codon-optimised *S. cerevisiae* Mcm5-R209E mutant was subcloned with *S. cerevisiae* wild type Mcm2, Mcm3, Mcm4, Mcm6, Mcm7, and Cdt1 in a pGB-dest backbone using GoldenBac assembly (*54*). The resulting MCM-Cdt1 co-expression plasmid was transformed into EMBacY cells for Bacmid preparation. Baculovirus preparation, insect cell expression and protein purification was carried out as previously described (*18*).

### Cloning, expression and purification of Dpb11

A synthetic gene encoding *S. cerevisiae* Dpb11, codon-optimised and fused to a C-terminal 3C protease site and 3xFLAG tag, was cloned into a pGB-04;05 shuttle vector. Following transformation into electro-competent EMBacY cells to generate bacmids, baculoviruses were transfected into Sf21 insect cells using FuGENE HD (Promega). Virus was amplified to P1; expression was performed in Sf21 cells insect cells (1 L at 1 million cells/mL) infected with 0.5% (v/v) P1 virus and harvested 48 hours after cell-cycle arrest.

Pelleted cells were resuspended in 50 mL Buffer A (25 mM HEPES-KOH pH 7.5, 500 mM KCl, 10% v/v glycerol, 0.02% w/v NP-40, 1 mM EDTA, 1 mM DTT) supplemented with 0.7 mM Phenylmethylsulphonylfluoride (PMSF) and a cOmplete EDTA-free protease inhibitor tablet (Merck). Lysis was achieved via sonication on ice (2 min total; 1 s on, 4 s off), followed by ultracentrifugation (45,000 rpm, 1 h, 4°C). The soluble fraction was treated with Bio-Lock Reagent (2.4 mL) and applied to a gravity column containing 1 mL anti-FLAG M2 Affinity Gel (Sigma). The column underwent washing with 150 mL Buffer A followed by 10 mL Buffer A containing 10 mM MgCl_2_ and 2 mM ATP. Protein elution was performed by incubating the resin with 5 mL Buffer A containing 0.5 mg/mL 3xFLAG peptide for 5 min. The eluate was adjusted to 150 mM KCl and purified via cation exchange on a 1 mL Mono S 5/50 column (Cytiva) in Buffer B (25 mM HEPES-KOH pH 7.5, 150 mM KCl, 10% v/v glycerol, 0.02% w/v NP-40, 1 mM EDTA, 1 mM DTT). Dpb11 was eluted using a 20-column volume linear gradient (150–1000 mM KCl). Peak fractions were pooled, dialysed overnight at 4°C against Buffer C (25 mM HEPES-KOH pH 7.5, 300 mM KOAc, 10% v/v glycerol, 0.02% NP-40, 1 mM EDTA, 1 mM DTT), concentrated to ∼0.5 mg/mL, and flash-frozen.

### Cloning, expression and purification of Sld2

The *S. cerevisiae* Sld2 sequence, tagged N-terminally with a retro-protein XXA solubility tag and C-terminally with a Twin-Strep-tag, was cloned into pET303 and transformed into *E. coli* T7 Express. Cultures (4 x 1 L LB with 100 µg/mL carbenicillin) were inoculated from colonies, incubated overnight at 37°C without shaking, then shifted to 30°C with agitation (200 rpm) until reaching an OD_600_ of 0.8. Expression was induced with 80 µM IPTG for 21 h at 30°C. Harvested cells were frozen at -80°C.

Upon thawing, cells were resuspended in 100 mL Buffer D (25 mM HEPES-KOH pH 7.5, 800 mM KCl, 10% v/v glycerol, 1 M Sorbitol, 2 mM ATP, 10 mM MgCl_2_, 0.02% v/v NP-40, 0.1% w/v Tween-20, 1 mM DTT) supplemented with protease inhibitors (2 cOmplete EDTA-free tablets and 0.7 mM PMSF). After sonication (2 min; 5 s on, 5 s off) and clarification (18,000 rpm, 20 min, 4°C), the lysate was loaded onto a 1 mL StrepTactinXT Superflow HighCapacity column. The resin was washed with 75 mL Buffer D followed by 25 mL Buffer E (25 mM HEPES-KOH pH 7.5, 500 mM NaCl, 10% v/v glycerol, 0.02% w/v NP-40, 1 mM EDTA, 1 mM DTT). Elution was achieved using Buffer E supplemented with Buffer BXT. Peak fractions were identified by SDS-PAGE, dialyzed into Buffer F (25 mM HEPES-KOH pH 7.5, 700 mM KOAc, 40% v/v glycerol, 0.02% w/v NP-40, 1 mM EDTA, 1 mM DTT), aliquoted at ∼0.8 mg/mL, and flash-frozen.

### Cloning, expression and purification of Twin-Strep-tagged Sumo-Mcm10

*S. cerevisiae* Mcm10 was engineered with an N-terminal 10xHis-SUMO tag and a C-terminal Twin-Strep-tag and expressed in *E. coli* Rosetta 2 pLysS. Dense overnight cultures were used to inoculate 6 x 1 L LB medium (containing carbenicillin and chloramphenicol). Cells were grown at 37°C to an OD_600_ of 0.7, induced with 0.5 mM IPTG, and cultured for 16 h at 16°C.

Pellets were lysed in 230 mL Buffer G (25 mM HEPES-KOH pH 7.6, 500 mM NaCl, 10% v/v glycerol, 1 mM EDTA, 0.05% w/v Tween-20, 1 mM DTT) with 4 cOmplete EDTA-free protease inhibitor tablets (Merck) via sonication (5 min; 2 s on, 5 s off). Following centrifugation (20,000 rpm in a Beckman Coulter JA-25.50 rotor, 30 min, 4°C), the supernatant was passed through a tandem setup of a 1 mL cOmplete His-Tag Purification column and a 1 mL StrepTactinXT 4Flow column. After washing with Buffer G, the protein was eluted from the His-column onto the Strep-column using Buffer G plus 200 mM imidazole. The His-column was removed, and the Strep-column was washed with Buffer B and eluted with Buffer H (Buffer G variant with 300 mM NaCl and 5 mM desthiobiotin). Mcm10 was dialyzed into Buffer I (25 mM HEPES pH 7.6, 200 mM NaCl, 20% v/v glycerol, 0.05% w/v Tween-20, 1 mM EDTA, 2 mM DTT), concentrated to ∼0.5 mg/mL, and frozen.

For the Mcm10 5D variant, aspartate substitutions were introduced at Phe113, Met114, Lys116, Phe117, and Lys121. For the Mcm10 3K variant, lysine substitutions were introduced at Phe401, Phe410, and Phe411. For the Mcm10 2E1A variant, the following amino acid substitutions were introduced: Phen340Glu, Arg347Ala, and Leu350Glu. All mutant proteins were expressed and purified using the same protocol as the wild-type, with the exception of using T7 express *LysY/l^q^*competent *E. coli* cells (NEB C3013) supplemented with 10 μg/ml carbenicillin.

### Assembly of MH-conjugated ARS1 Templates

A 168-bp DNA fragment comprising the *S. cerevisiae* ARS1 origin flanked by MH recognition sites was generated via PCR. This template was covalently coupled to tandem ALFA/Twin-Strep-tagged MH using established conjugation protocols (*14*).

### dCMGE Assembly and Activation in a test tube

Complex assembly and activation followed published methods (*15, 16*) with adjustments. MCM-DHs were loaded by incubating 40 nM MH-ARS1, 104 nM ORC, 104 nM Cdc6, and 285 nM Mcm2-7/Cdt1 in 100 µL Buffer J (25 mM HEPES-KOH pH 7.5, 100 mM potassium L-glutamate, 10 mM Mg(OAc)2, 1 mM DTT, 1 mM ATP, 0.02% w/v NP-40) for 45 min at 30°C.

For Mcm2-7/Cdt1 RA mutants, concentrations were adjusted to 382 nM Mcm2-7/Cdt1(Mcm3RA), 724 nM Mcm2-7/Cdt1(Mcm7RA), or 300 nM Mcm2-7/Cdt1(Mcm4RA). Complexes were phosphorylated by adding 118 nM DDK (10 min, 24°C) and immobilised on 50 µL MagStrep “type3” XT beads in Buffer K (Buffer J variant without ATP). Following 3 x 200 µL washes with Buffer L (25 mM HEPES-KOH pH 7.5, 500 mM NaCl, 5 mM Mg(OAc)_2_, 0.02% w/v NP-40) and 1x 200 µL wash with Buffer K, complexes were eluted with 20 mM biotin in 100 µL Buffer J. The eluted MCM-DHs were split into five 20 µL reactions and dCMGE formation and activation was initiated by adding 200 nM S-CDK, 75 nM Dpb11, 63 nM Polε, 252 nM GINS, 221 nM Cdc45, 75 nM Sld3/7, 126 nM Sld2, and 50 nM Mcm10 (wild-type or mutant) to each 20 µL reaction. Reactions proceeded for 12 min at 30°C and were immobilized on 15 µL ALFA PE Selector slurry for 5 min at 24°C. Following 3 x 200 µL washes with Buffer K, complexes were eluted with 200 µM ALFA peptide in 20 µL Buffer J for 10 min at 24°C. Subsequently, samples were subjected to nsEM or cryo-EM analysys (detailed below).

### *In Vitro* DNA Replication

Replication assays were performed at 30°C and 1250 rpm following previous protocols (*55*). The MCM-DH loading reaction (20 min, 30°C) contained 40 nM ORC, 40 nM Cdc6, 60 nM Mcm2-7/Cdt1, and 4 nM pJY22 plasmid DNA (10.6 kb) in Buffer M (25 mM HEPES-KOH pH 7.6, 100 mM potassium L-glutamate, 10 mM Mg(OAc)_2_, 5 mM ATP, 0.02% NP-40-S and 2 mM DTT).

Loaded MCMs were treated with 50 nM DDK (15 min). Replication was initiated by adding firing factors (40 nM Dpb11, 20 nM Pol epsilon, 20 nM GINS, 80 nM Cdc45, 20 nM CDK, 25 nM Sld3/7, 50 nM Sld2, 200 nM RPA, 20 nM TopoI, 50 nM Pol alpha, 20 nM Mcm10) and nucleotides (200 μM each CTP/GTP/UTP, 80 μM each dNTP and 33 nM ɑ^32^P-dCTP). Where indicated, Mcm10 and Mcm2-7/Cdt1 wild-type proteins were replaced by mutants. Reactions were quenched with 85 mM EDTA, purified with an Illustra MicroSpin G-50 column, denatured in 2% sucrose/0.02% bromophenol blue/60 mM NaOH/10 mM EDTA, resolved on alkaline 0.8% agarose gels containing 30 mM NaOH and 2 mM EDTA at 1 V/cm for 17h, fixed in cold 5% trichloroacetic acid, dried, exposed to phosphor screens, and scanned with a Typhoon phosphor imager.

### nsEM Sample Preparation, Data Acquisition and Analysis

Samples (4 µL) were applied for 2 min to glow-discharged (25 mA, 1 min in a Quorum GloQube Plus) carbon-coated 400-mesh copper grids (Agar), stained with two applications of 4 µL 2% uranyl acetate for 45 s, blotted dry and imaged on a FEI Tecnai G2 Spirit Twin (120 kV) with a Gatan Rio16 camera. Between 50 and 150 micrographs were collected for each experiment and per replicate. Data were processed using Relion 5.0 (*56*), Topaz (*57*) and cryoSPARC v4.7.1 (*58*). Particles were 2D classified in cryoSPARC (*58*) to identify CMGE states (dCMGE, dCMGE-Mcm10, sCMGE) and quantified as the ratio of specific complexes to total CMGE-containing species. Mcm10-dependent dCMGE splitting efficiency was calculated as number of sCMGE particles multiplied by a factor of 0.5 (to account for the fact that two sCMGEs originate from a single replication origin) divided by the number of all replication origins where dCMGE assembly had occured.

### Cryo-EM Sample Preparation of dCMGE10^Mcm7RA^

Double CMGE(Mcm7RA)-Mcm10 complexes were prepared as described above (*In vitro* dCMGE Assembly and Activation) with the exception that the elution buffer from the ALFA PE Selector slurry contained 1 mM ATPγS instead of 1 mM ATP. UltrAuFoil R1.2/1.3 300 mesh (Quantifoil) cryo-EM grids were glow discharged (40 mA, 5 min in a Quorum GloQube Plus) and incubated twice for 2 min with 4 µL of 0.22 mg/mL graphene oxide suspension in water. Grids were washed with two 20 µL droplets of water from the frontside, blotted and washed with a third droplet from the backside. Four applications of 4 µL sample were applied for 2 min to each grid in a Mark IV Vitrobot (FEI) at 24°C and 90% humidity. Grids were blotted for 4.5 s at blotforce 0, plunge-frozen in liquid ethane and stored in liquid nitrogen.

### Cryo-EM Sample Preparation of sCMGE ^Mcm4RA^

Single CMGE(Mcm4RA) complexes were prepared as described above for dCMGE(Mcm7RA)-Mcm10, with the exception that six applications of 4 µL sample were applied to each cryo-EM grid instead of four during vitrification.

### Cryo-EM Data Acquisition of dCMGE10 ^Mcm7RA^

Data were collected on a 300kV FEI Titan Krios G3i equipped with a Falcon 4 detector in counting mode and Selectris energy filter (10eV slit). 178,563 movies were recorded in EER format at 130,000x magnification (0.95 Å/px), with spot size 9, 660 nm beam diameter, a 100 µm objective aperture and a defocus range from -2.0 to -2.9 µm with a total fluence of 42.8 electrons/Å^2^.

### Cryo-EM Data Acquisition of sCMGE ^Mcm4RA^

Data were collected on a 300kV FEI Titan Krios G3i equipped with a Falcon 4 detector in counting mode and Selectris energy filter (10eV slit). 138,014 movies were recorded in LZW-compressed TIFF format at 130,000x magnification (0.95 Å/px), with spot size 9, 660 nm beam diameter, a 100 µm objective aperture and a defocus range from -1.6 to -2.2 µm with a total fluence of 52.7 electrons/Å^2^.

### Cryo-EM Image Processing of dCMGE10 ^Mcm7RA^

178,563 EER movies were motion-corrected using Relion 5.0 (*56*)’s own implementation of MOTIONCOR2 (*59*) and CTF-estimated with CTFFind 4.1 (*60*). 576 particles were manually picked from 30 micrographs and used as input for Topaz (*57*) training in Relion 5.0 (*56*) (400 Å particle diameter, 50 particles expected per micrograph). The trained Topaz model was used to pick a subset of 10,000 micrographs. 358,585 particles were extracted at 3.8 Å/px (4x binning) with a 128 px box and cleaned up with multiple rounds of 2D Classification in cryoSPARC v4.7.1 (*58*). A subset of clean dCMGE10, dCMGE, and MCM-DH classes were selected to generate initial 3D models using cryoSPARC’s ab initio reconstruction (*58*). The refined particle stack was used to train a second Topaz (*57*) model in Relion 5.0 (*56*) and pick a second subset of 10,000 micrographs. This process was iteratively repeated 10 times, resulting in 10 different Topaz (*57*) models. The full dataset of 178,563 micrographs was picked using all 10 Topaz (*57*) models in Relion 5.0 (*56*), generating 10 particle stacks of ∼ 4,000,000 particles each extracted at 3.8 Å/px (4x binning) with a 128 px box. Each particle stack was cleaned up with multiple rounds of 2D Classification in cryoSPARC (*58*) followed by multiple rounds of Heterogenous Refinement using the dCMGE10, dCMGE and MCM-DH classes from the ab initio reconstruction as well as two junk classes. Particles from the dCMGE10 class were merged, duplicates removed (distance cutoff for removal 100 Å), and the resulting 530,710 particles were subjected to multiple rounds of Heterogenous Refinement. The final stack of 349,679 particles was unbinned to 0.95 Å/px with a 512 px box, refined using cryoSPARC’s Non-Uniform Refinement (*58*) with C2 symmetry applied, and symmetry expanded to yield a stack of 699,358 particles. To recover symmetry breaking features, the C2 symmetry-expanded dCMGE10 particles were 3D classified without alignment in cryoSPARC (*58*) into 10 classes, limiting the resolution to 10 Å. Two classes of 67,110 (State II) and 76,228 (State IV) particles that showed high resolution features for both CMGs in the dimer and different Pol epsilon engagement states were 3D refined in C1 with local searches and Blush (*61*) in Relion 5.0 (*56*), subjected to CTF refinement and Bayesian Polishing (*62*), and 3D refined in C1 with local searches and Blush (*61*) to a nominal resolution of 3.52 Å (State II) and 3.92 Å (State IV). To overcome the loss of structural features caused by flexibility of the dCMGE10 complex, Dynamight (*63*) flexibility analysis was performed on the two particle stacks. Motion estimation was done using 30,000 gaussians, inverse deformations were estimated over 200 epochs, and after deformed backprojection the resulting half maps were post-processed without sharpening (B-factor 0) to a final resolution of 3.42 Å (State II) and 3.47 Å (State IV). To aid interpretation and model building, the two half maps generated from Dynamight (*63*) analysis were used as input for LocScale-2.0 (*64*) confidence-weighted density modification (LocScale-FEM), as well as for EMReady2 (*65*) local quality-aware density modification.

To distinguish all configurations a single CMGE could adopt in the Mcm10-locked dimer, the symmetry-expanded stack of 699,358 particles was subjected to masked Local Refinement in cryoSPARC (*58*) masking a single CMGE-Mcm10, followed by masked 3D classification without alignment into 5 classes. Apart from the previously observed CMGE configurations of State II (Pol epsilon engaged to CMG) and State IV (Pol epsilon disengaged from CMG), a third class of 82,776 particles showed Pol epsilon partially disengaged from CMG. This particle stack was subjected to masked 3D refinement in C1 with local searches and Blush (*61*) in Relion 5.0 (*56*), two rounds of CTF refinement, one round of Bayesian Polishing (*62*), and masked 3D refinement in C1 with local searches and Blush (*61*) to a nominal resolution of 3.38 Å. To recover the signal of the CMGE that had been masked out, the particles were 3D refined in C1 with local searches and Blush (*61*) without a mask to a nominal resolution of 3.53 Å, and 3D classified without alignment into 4 classes using a mask on the poorer-quality CMGE and limiting resolution to 10 Å. Two classes of 23,793 (State I) and 27,138 (State III) particles that showed high resolution features for the masked CMG in the dimer and different Pol epsilon engagement states were 3D refined in C1 with local searches and Blush (*61*) without a mask to a nominal resolution of 4.05 Å (State I) and 3.92 Å (State III). Dynamight (*63*) flexibility analysis was performed on the two particle stacks as described above for State II and State IV. The resulting half maps were post-processed without sharpening (B-factor 0) to a final resolution of 3.42 Å (State I) and 3.47 Å (State III). To aid interpretation and model building, the two half maps generated from Dynamight (*63*) analysis were used as input for LocScale-2.0 (*64*) confidence-weighted density modification (LocScale feature-enhanced map), as well as for EMReady2 (*65*) local quality-aware density modification.

### Cryo-EM Image Processing of sCMGE ^Mcm4RA^

138,014 LZW-compressed TIFF movies were motion-corrected using Relion 5.0 (*56*)’s own implementation of MOTIONCOR2 (*59*) and CTF-estimated with CTFFind 4.1 (*60*). 583 particles were manually picked from 20 micrographs and used as input for Topaz (*57*) training in Relion 5.0 (*56*) (400 Å particle diameter, 70 particles expected per micrograph). The trained Topaz model was used to pick a subset of 10,000 micrographs. 629,304 particles were extracted at 3.8 Å/px (4x binning) with a 128 px box and cleaned up with multiple rounds of 2D Classification in cryoSPARC v4.7.1 (*58*). A subset of clean sCMGE, dCMGE10, dCMGE, and MCM-DH classes were selected to generate initial 3D models using cryoSPARC’s ab initio reconstruction (*58*). The refined particle stack was used to train a second Topaz (*57*) model in Relion 5.0 (*56*) and pick the full dataset. 8,288,404 particles were extracted at 3.8 Å/px (4x binning) with a 128 px box and cleaned up with multiple rounds of 2D Classification in cryoSPARC (*58*) followed by multiple rounds of Heterogenous Refinement using the sCMGE, dCMGE10, dCMGE and MCM-DH classes from the ab initio reconstruction as well as two junk classes. 1,053,784 particles from the sCMGE class were refined using cryoSPARC’s Non-Uniform Refinement (*58*) in C1 to a nominal resolution of 7.91 Å (Nyquist after binning 7.6 Å). The refined particles were 3D classified without alignment in cryoSPARC (*58*) into 7 classes, limiting the resolution to 10 Å. One class of 169,397 particles that showed density for the ejected lagging strand template was further 3D classified without alignment in cryoSPARC (*58*) into 8 classes, limiting the resolution to 10 Å. The 37,837 particles from the class with the best continuous density for the ejected lagging strand template were unbinned to 0.95 Å/px with a 420 px box, refined in C1 with local searches and Blush (*61*) in Relion 5.0 (*56*), subjected to two rounds of CTF refinement and Bayesian Polishing (*62*), and 3D refined in C1 with local searches and Blush to a nominal resolution of 3.09 Å. To overcome the loss of structural features caused by flexibility of the ejected lagging strand, Dynamight (*63*) flexibility analysis was performed on the particle stack. Motion estimation was done using 30,000 gaussians, inverse deformations were estimated over 200 epochs, and after deformed backprojection the resulting half maps were post-processed without sharpening (B-factor 0) to a final resolution of 3.05 Å. To aid interpretation and model building, the two half maps generated from Dynamight (*63*) analysis were used as input for LocScale-2.0 (*64*) confidence-weighted density modification (LocScale feature-enhanced map).

### Atomic Model Building and Refinement of dCMGE10 ^Mcm7RA^

Models were built into locally-refined maps density-modified with Locscale-2.0 (*64*) feature-enhanced map. First, a single CMGE-Mcm10 complex without DNA was built into the highest resolution State II map. A single CMGE complex from PDB entry 7PMK (*66*) was fitted as a rigid body into the density using UCSF ChimeraX v1.10.1 (*67*). Then, the N-terminal and ATPase tiers of the Mcm2-7 subunits, and each individual chain of GINS, Cdc45 and Pol epsilon were fitted as separate rigid bodies. As it strikingly matched the cryo-EM density, an AlphaFold3 (*68*) structure prediction of Mcm10-Cdc45-Mcm2-Mcm6 was docked as a rigid body into the density, Cdc45-Mcm2-Mcm6 were deleted and the remaining Mcm10 chain was used as a starting model for Mcm10. Each chain of the complex was subject to three consecutive rounds of flexible fitting in Coot v0.9.8.1 (*69*) with three different chain self-restraints (rigid 5, intermediate 4.3, weak 3.7). Fragments not supported by the density were deleted. Five ATPase sites were identified as occupied, with ATPγS modelled based on the density (Mcm4/6, 6/2, 2/5, 5/3 and 3/7). An alanine residue was introduced at position Mcm7 R593. Subsequently, density fit and side-chain rotamers were real-space refined manually in Coot (*69*) for all chains. Finally, the entire model was subjected to one round of molecular-dynamics flexible fitting in ISOLDE v1.763 (*70*) to reduce clashes.

Following the same procedure, but with the newly built single CMGE-Mcm10 complex as starting model, a single CMG-Mcm10 complex (with Pol epsilon disengaged) was built into State IV map. All six ATPase sites were identified as occupied, with ATPγS modelled based on the density.

To complete the protein model for State II and State IV, a single CMG-Mcm10 monomer was rigid-body docked into the symmetry-equivalent density corresponding to the second sCMG using ChimeraX (*67*). Each chain of the complex was flexibly fitted using Coot (*69*) as described above, density fit and side-chain rotamers were real-space refined manually, and clashes were reduced via molecular-dynamics flexible fitting in ISOLDE (*70*).

To build the protein models for State I and State III, two sCMGE-Mcm10 models (State I) or one sCMGE-Mcm10 model and one sCMG-Mcm10 model (State III) refined to State II map were rigid-body docked into their respective cryo-EM maps using ChimeraX (*67*). The N-terminal and ATPase tiers of the Mcm2-7 subunits, and each individual chain of GINS, Cdc45, Pol epsilon and Mcm10 were fitted as separate rigid bodies. Each chain of the complex was flexibly fitted in Coot (*69*) as described above. Five ATPase sites were identified as occupied in each CMG of State I, with ATPγS (Mcm4/6, 2/5, 5/3 and 3/7) or ADP (Mcm6/2) modelled based on the density. In the map of State III, five ATPase sites were identified as occupied by ATPγS in the CMG with visible Pol epsilon density (Mcm4/6, 6/2, 2/5, 5/3 and 3/7), whereas all six ATPase sites were identified as occupied by ATPγS in the CMG without visible Pol epsilon density. For both State I and State III models, density fit and side-chain rotamers were real-space refined manually, followed by molecular-dynamics flexible fitting in ISOLDE (*70*) to reduce clashes.

DNA was built last for all models. The DNA from PDB 7Z13 (dCMGE) (*15*) was first rigid-body docked into State I map using ChimeraX (*67*). It was then flexibly fitted in Coot (*69*), positioning the backbone phosphates of the leading strand template in contact with the PS1 beta-hairpin lysines of Mcm2, Mcm5, Mcm3, and Mcm7 of both CMGs accounting for the symmetric 2 bp advancement via Mcm7. A stretch of B-form DNA at the dimerization interface that sterically clashed with the Mcm2 Zinc Fingers was deleted and nucleotides were manually rebuilt inside the density as an open bubble using Coot (*69*).

To build the DNA for State II, DNA from State I was first rigid-body docked into the map of State II in ChimeraX (*67*). It was then flexibly fitted using Coot (*69*) and split in two halves. The half of the DNA where CMG contacts persisted unchanged from State I was not modified, whereas the opposite half of the DNA was positioned in the density so that the backbone phosphates of the leading strand template were in contact with the PS1 beta-hairpin lysines of Mcm7, Mcm4, Mcm6, and Mcm2, accounting for the 6 bp advancement of the CMG via Mcm4, Mcm6, and Mcm2. The resulting stretching of the DNA was modelled by re-joining the two halves of the DNA together and real-space refining the region between the backbone phosphate of Adenosine 34, contacted by the Mcm2 PS1 beta-hairpin lysine (K633), and the open bubble at the dimerization interface.

To build the DNA for State III, DNA from State II was first rigid-body docked into the map of State III using ChimeraX (*67*) and then subjected to three consecutive rounds of flexible fitting in Coot (*69*) with decreasingly stringent molecule self-restraints.

To build the DNA for State IV, DNA from State III was rigid-body docked into State IV map in ChimeraX (*67*), flexibly fitted in Coot (*69*) and split in two. The half of the DNA where CMG contacts persisted (unchanged from State III) was not modified, whereas the opposite half was flexibly fitted in the density so that the backbone phosphates of the leading strand template were in contact with the PS1 beta-hairpin lysines of Mcm7, Mcm4, Mcm6, and Mcm2, accounting for the 6 bp advancement of the CMG via Mcm4, Mcm6, and Mcm2. The resulting stretched DNA was modelled by joining the two halves of the DNA together and real-space refining the region between the backbone phosphate of Adenosine 34, contacted by the Mcm2 PS1 beta-hairpin lysine (K633), and the melted bubble at the dimerization interface.

After building the DNA for each model, a final round of molecular-dynamics flexible fitting in ISOLDE (*70*) was performed to reduce clashes, followed by real-space refinement in Phenix v1.21.1 (*71*) using default settings against the respective unmodified maps. For all structures, the quality of the resulting atomic models was evaluated with MolProbity (*72*) (Table S1).

### Atomic Model Building and Refinement of sCMGE ^Mcm4RA^

The model was built into a locally-refined map density-modified with Locscale-2.0 (*64*) feature-enhanced map. A single CMGE complex from PDB entry 7PMK (*66*) was fitted as a rigid body into the density using UCSF ChimeraX v1.10.1 (*67*). Then, the N-terminal and ATPase tiers of the Mcm2-7 subunits, and each individual chain of GINS, Cdc45 and Pol epsilon were fitted as separate rigid bodies. An AlphaFold3 (*68*) structure prediction of *Haemophilus parainfluenzae* Type II methyltransferase bound to a 13-bp-long DNA duplex was docked as a rigid body into the density and used as a starting model for the methyltransferase. PDB 8T1U was used as the starting model for the methyltranferase-bound DNA containing a 5-fluoro,5-methyl-2’-deoxy-cytidine. An 8-bp-long B-form DNA duplex was generated in Coot v0.9.8.1 (*69*) and fitted as a rigid body into the C-terminal duplex DNA density. Each chain of the complex was subjected to three consecutive rounds of flexible fitting in Coot (*69*) using decreasingly stringent chain self-restraints (rigid 5, intermediate 4.3, weak 3.7). Fragments not supported by the density were deleted. All six ATPase sites were identified as occupied, with ATPγS modelled based on the density. An alanine residue was introduced at position Mcm4 R701. Subsequently, density fit and side-chain rotamers were real-space refined manually in Coot (*69*) for all chains. Finally, the entire model was subjected to one round of molecular-dynamics flexible fitting in ISOLDE v1.763 (*70*) to reduce clashes followed by real-space refinement in Phenix v1.21.1 (*71*) against the unmodified map with restraints for 5-fluoro,5-methyl-2’-deoxy-cytidine generated using the eLBOW tool (*73*). The quality of the resulting atomic model was evaluated with MolProbity (*72*) (Table S1).

### Energy-minimised morph movie generation with RigiMOL in PyMol

To create a molecular movie transitioning from dCMGE (*15*) (PDB 7Z13) through State I, State II, State III to State IV, morph movies between pairs of states (dCMGE to I, I to II, II to III, III to IV) were generated first. To ensure that the models to morph were aligned and had the same number of atoms, labelled the same, the model pairs were opened in ChimeraX v1.10.1 (*67*) as #1 and #2, aligned on their Cdc45 subunits with the ChimeraX (*67*) matchmaker and the following command was executed via the command line: “morph #1,2 same true”. The pdb files of the first and last frames of the ChimeraX (*67*) morph were saved, opened in PyMol v3.1.5.1 and from the first pdb file’s object menu panel the following options were selected: A > generate > morph > wizard. A RigiMOL morph was thus created with 30 steps and 10 refinement cycles, and its frames were saved as pdb files, opened in ChimeraX (*67*) for visualization purposes and the following command was executed via the command line: “morph #1.1-30 same true frames 1”.

## Acknowledgments

We would like to thank members of the Costa lab for useful discussion, the Crick Structural Biology STP for computational support (A. Purkiss and A. Nans); yeast cultures (N. Patel, A. Alidoust and D. Patel) and cryo-EM support (A. Nans, R. Baradaran and N. Lukoyanova).

## Author contributions

G.P. and A.C. conceived the study and wrote the manuscript with input from all authors. G.P. performed all experiments with the following exceptions. B.C. and M.H.G. performed DNA replication assays under the supervision J.F.X.D. T.P. and A.B. cloned and purified certain protein variants. A.B. contributed to atomic model building. A.C. supervised the study.

## Data availability

Cryo-EM density maps have been deposited in the Electron Microscopy Data Bank (EMDB) under the following accession codes: EMD-57126 (maps for the dCMGE10^Mcm7RA^ State I), EMD-57127 (maps for the dCMGE10^Mcm7RA^ State II), EMD-57128 (maps for the dCMGE10^Mcm7RA^ State III), EMD-57130 (maps for the dCMGE10^Mcm7RA^ State IV) and EMD-57131 (maps for the sCMGE^Mcm4RA^). Atomic coordinates have been deposited in the Protein Data Bank (PDB) with the following accession codes: 29ER (dCMGE10^Mcm7RA^ State I), 29ES (dCMGE10^Mcm7RA^ State II), 29ET (dCMGE10^Mcm7RA^ State III), 29EV (dCMGE10^Mcm7RA^ State IV) and 29EW (sCMGE^Mcm4RA^).

## Supplementary Figures

**Fig. S1.**
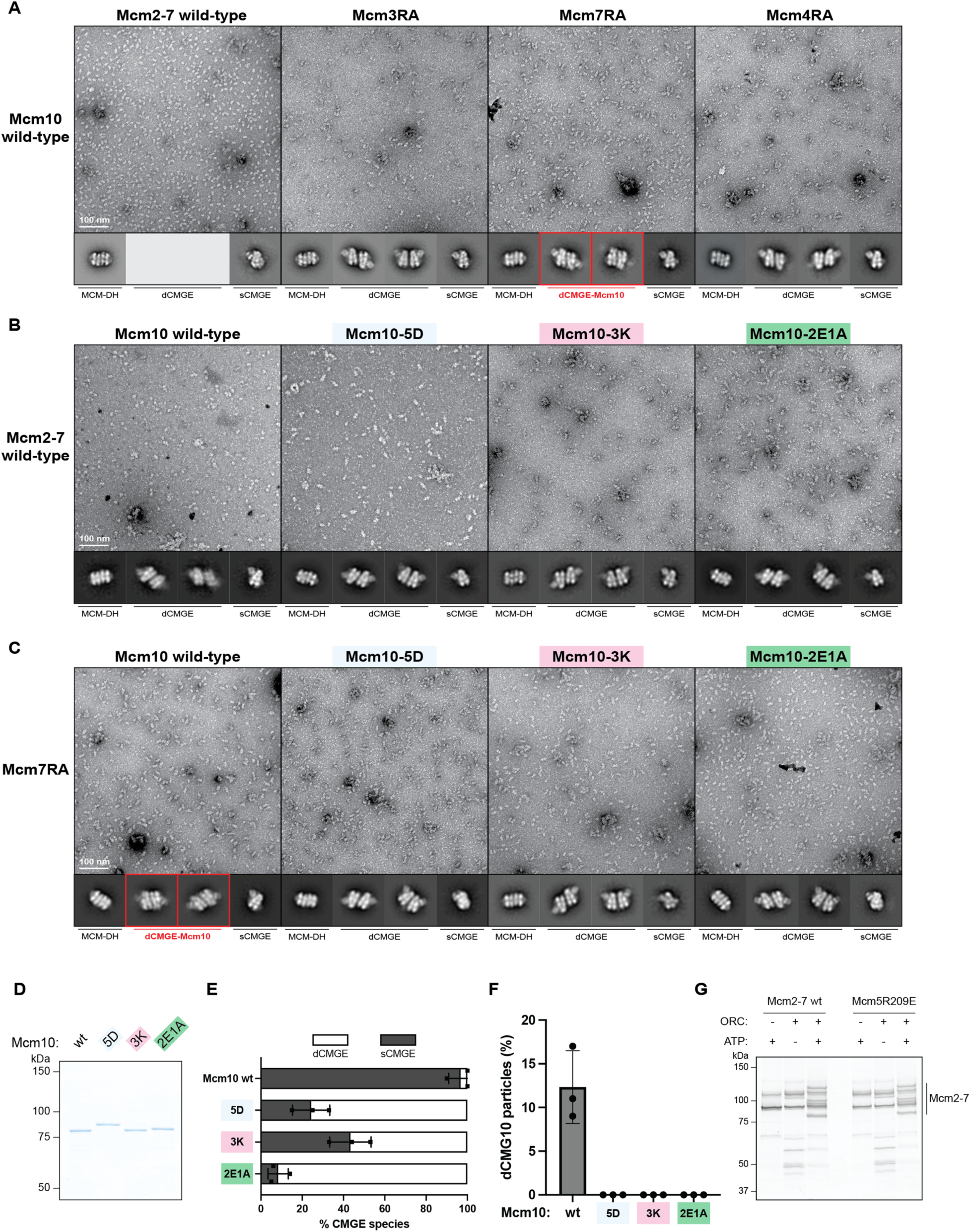
(**A**) Representative negative-stain EM micrographs and 2D averages of CMG activation reactions carried out with Mcm10 wild type and Mcm2-7 wild type, Mcm3RA, Mcm7RA or Mcm4RA. Red boxes mark unique dCMGE-Mcm10 averages obtained with Mcm7RA where N-terminal MCMs were tightly interacting. (**B**) Representative negative-stain EM micrographs and 2D averages of CMG activation reactions carried out with Mcm2-7 wild type and Mcm10 wild type, Mcm10-5D, Mcm10-3K or Mcm10-2E1A. (**C**) Representative negative-stain EM micrographs and 2D averages of CMG activation reactions carried out with Mcm7RA and Mcm10 wild type, Mcm10-5D, Mcm10-3K or Mcm10-2E1A. Red boxes mark dCMGE-Mcm10 averages obtained with Mcm10 wild type but not with the mutants. (**D**) Purified Mcm10 wild type, Mcm10-5D, Mcm10-3K or Mcm10-2E1A analysed by SDS–PAGE with Coomassie staining. (**E**) Bar graph comparing CMGE species in reactions carried out with Mcm2-7 wild type and Mcm10 wild type, Mcm10-5D, Mcm10-3K or Mcm10-2E1A, as shown in **b**. This experiment was performed three times. (**F**) Bar graph comparing dCMG10 complex formation in reactions carried out with Mcm7RA and Mcm10 wild type, Mcm10-5D, Mcm10-3K or Mcm10-2E1A, as shown in **c**. This experiment was performed three times. (**G**) Silver stain gel of MCM double hexamer loading reactions carried out with Mcm2-7 wild type or Mcm5R209E.

**Fig. S2.**
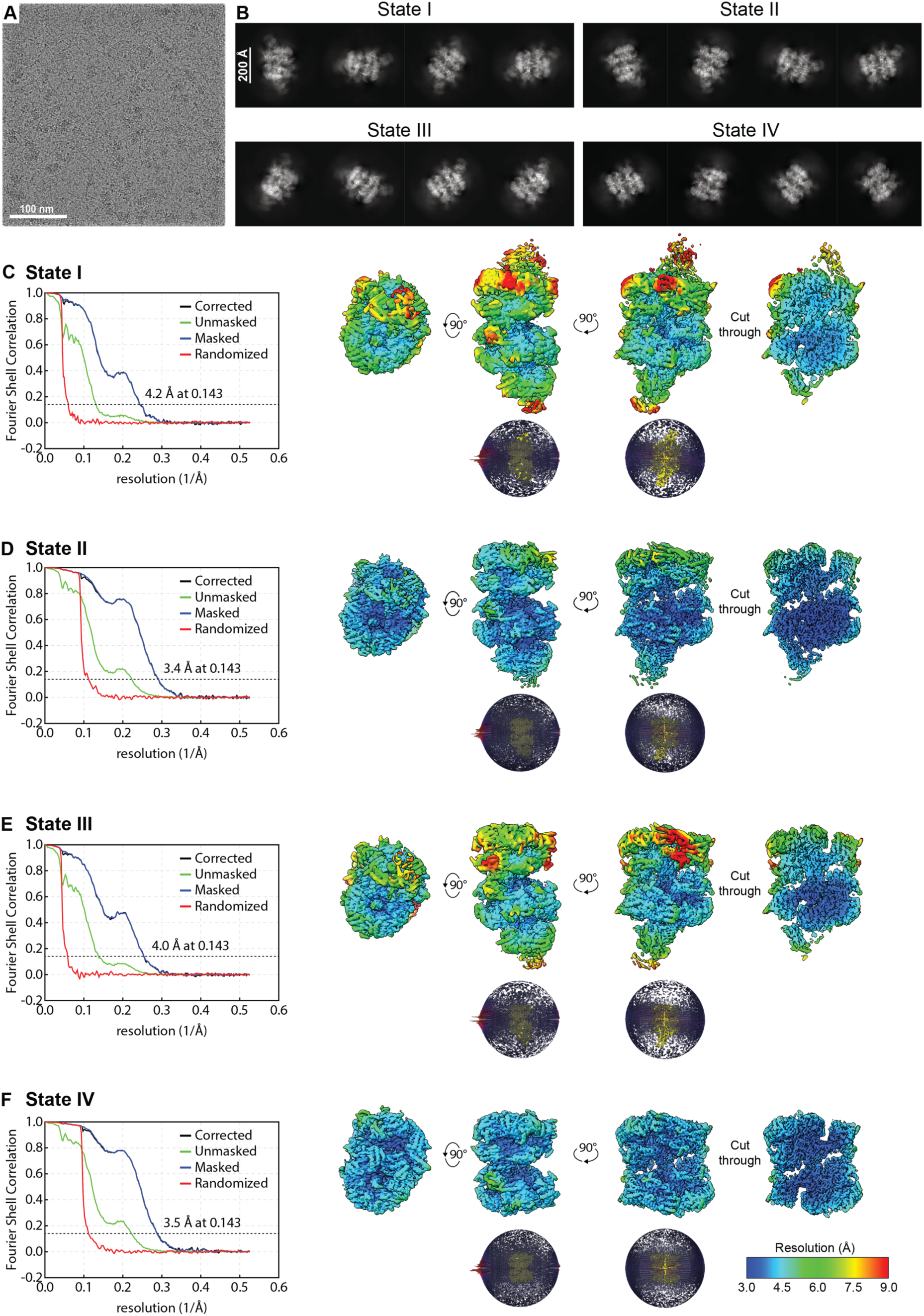
(**A**) Representative cryo-electron micrograph from the dCMGE10^Mcm7RA^ dataset. (**B**) Cryo-EM 2D averages of dCMGE10^Mcm7RA^ State I, State II, State III and State IV. (**C**) Fourier shell correlation plot, four views of the map coloured according to the local resolution, and angular distribution of dCMGE10^Mcm7RA^ State I. (**D**) Fourier shell correlation plot, four views of the map coloured according to the local resolution, and angular distribution of dCMGE10^Mcm7RA^ State II. (**E**) Fourier shell correlation plot, four views of the map coloured according to the local resolution, and angular distribution of dCMGE10^Mcm7RA^ State III. (**F**) Fourier shell correlation plot, four views of the map coloured according to the local resolution, and angular distribution of dCMGE10^Mcm7RA^ State IV.

**Fig. S3.**
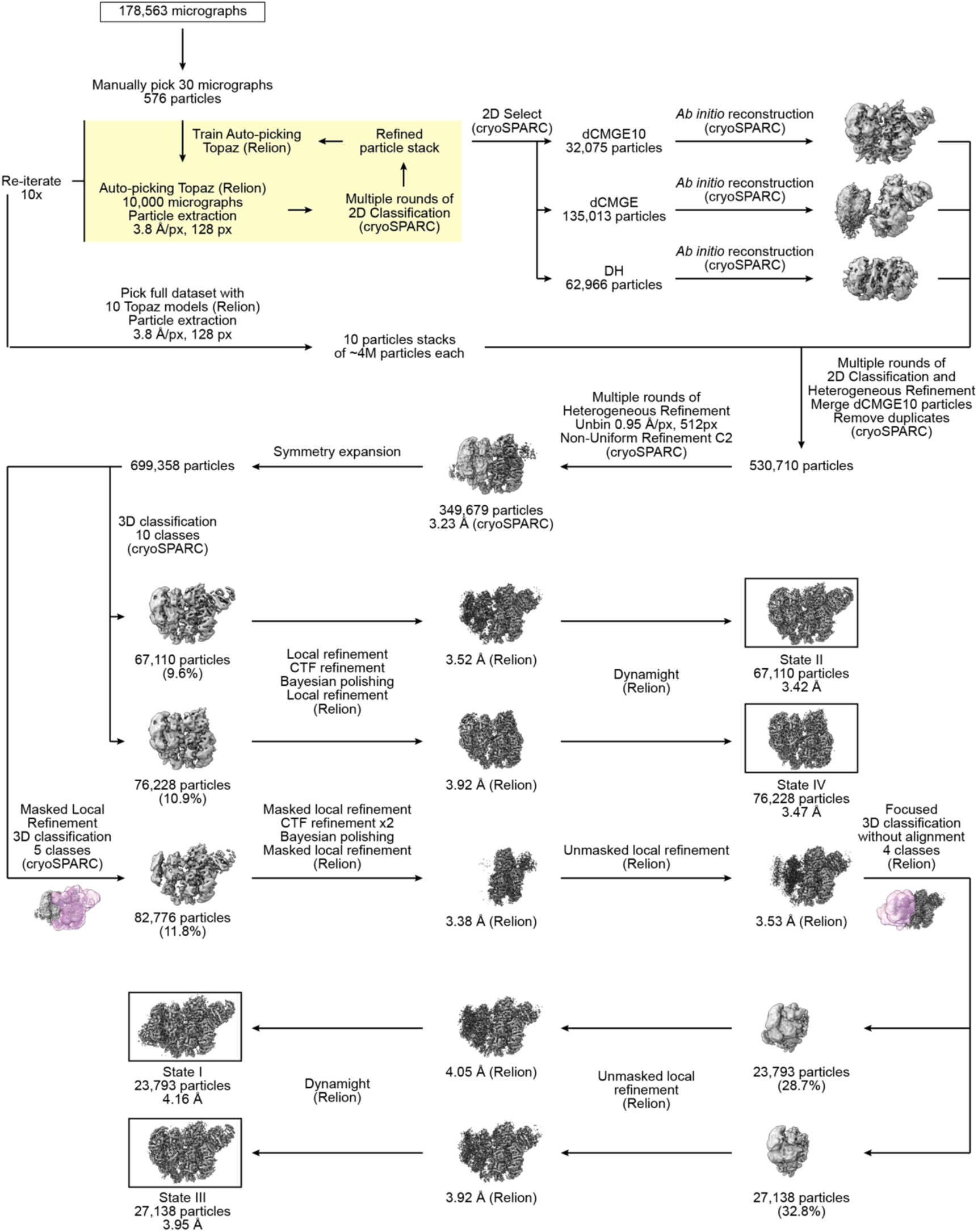
Image processing pipeline for dCMGE10^Mcm7RA^ State I, State II, State III, and State IV.

**Fig. S4.**
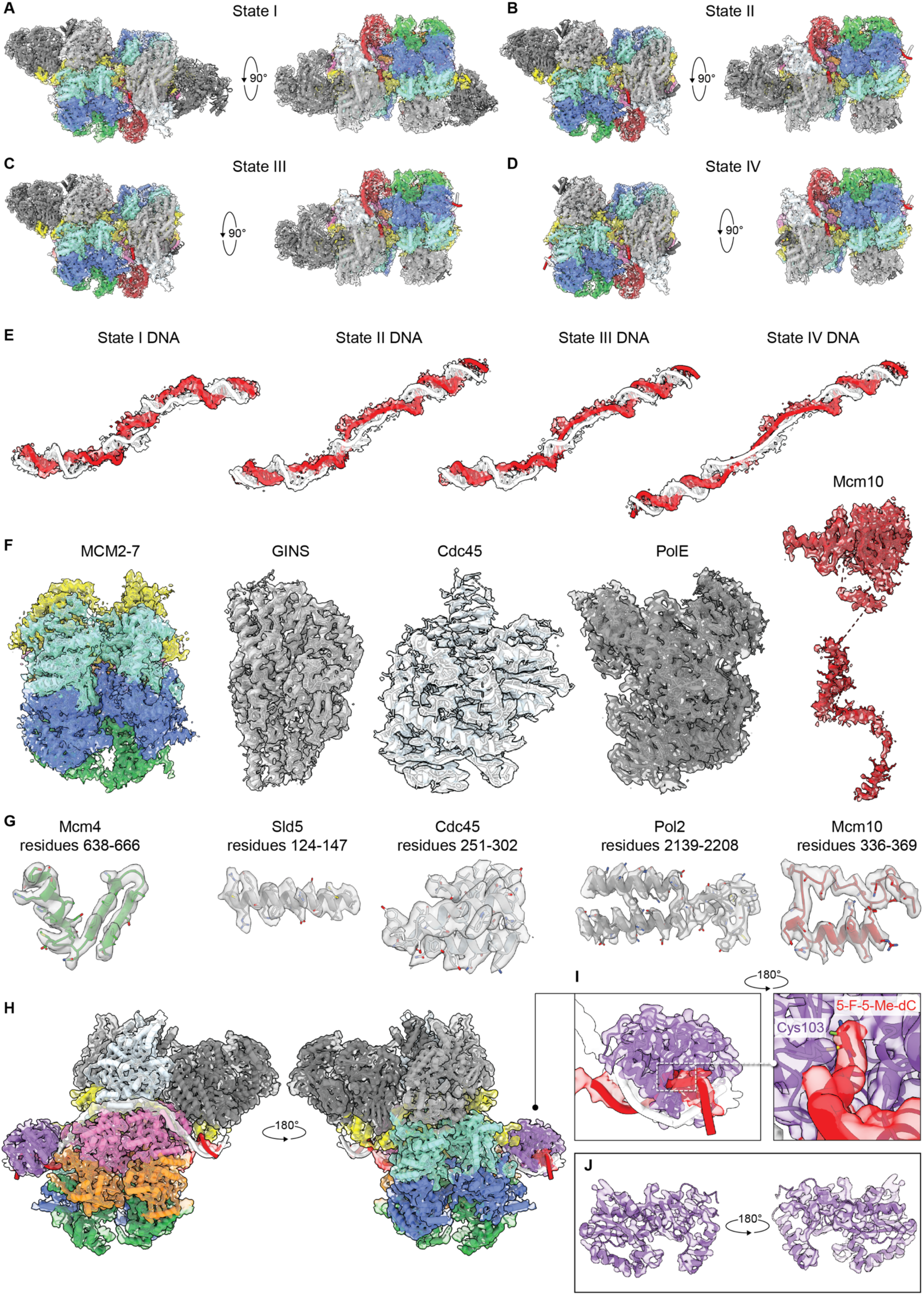
(**A**) Cryo-EM map of dCMGE10^Mcm7RA^ State I with fitted model. (**B**) Cryo-EM map of dCMGE10^Mcm7RA^ State II with fitted model. (**C**) Cryo-EM map of dCMGE10^Mcm7RA^ State III with fitted model. (**D**) Cryo-EM map of dCMGE10^Mcm7RA^ State IV with fitted model. (**E**) Cryo-EM maps of DNA from dCMGE10^Mcm7RA^ State I through IV with fitted models. (**F**) Cryo-EM maps of MCM2-7, GINS, Cdc45, DNA polymerase epsilon, and Mcm10 with fitted models. (**G**) Cryo-EM maps with fitted models of Mcm4 residues 638-666, Sld5 residues 124-147, Cdc45 residues 251-302, Pol2 residues 2139-2208, and Mcm10 residues 336-369. (**H**) Cryo-EM map of sCMGE^Mcm4RA^ with fitted model. (**I**) First inset shows the Cryo-EM map with fitted model of sCMGE^Mcm4RA^ DNA bound to MH methyltransferase. The second inset shows a zoomed-in view of the flipped-out 5-fluoro,5-methyl-2’-deoxy-cytidine covalently linked to MH Cys103. (**J**) Cut-through view of Cryo-EM map of MH methyltransferase with fitted model.

**Fig. S5.**
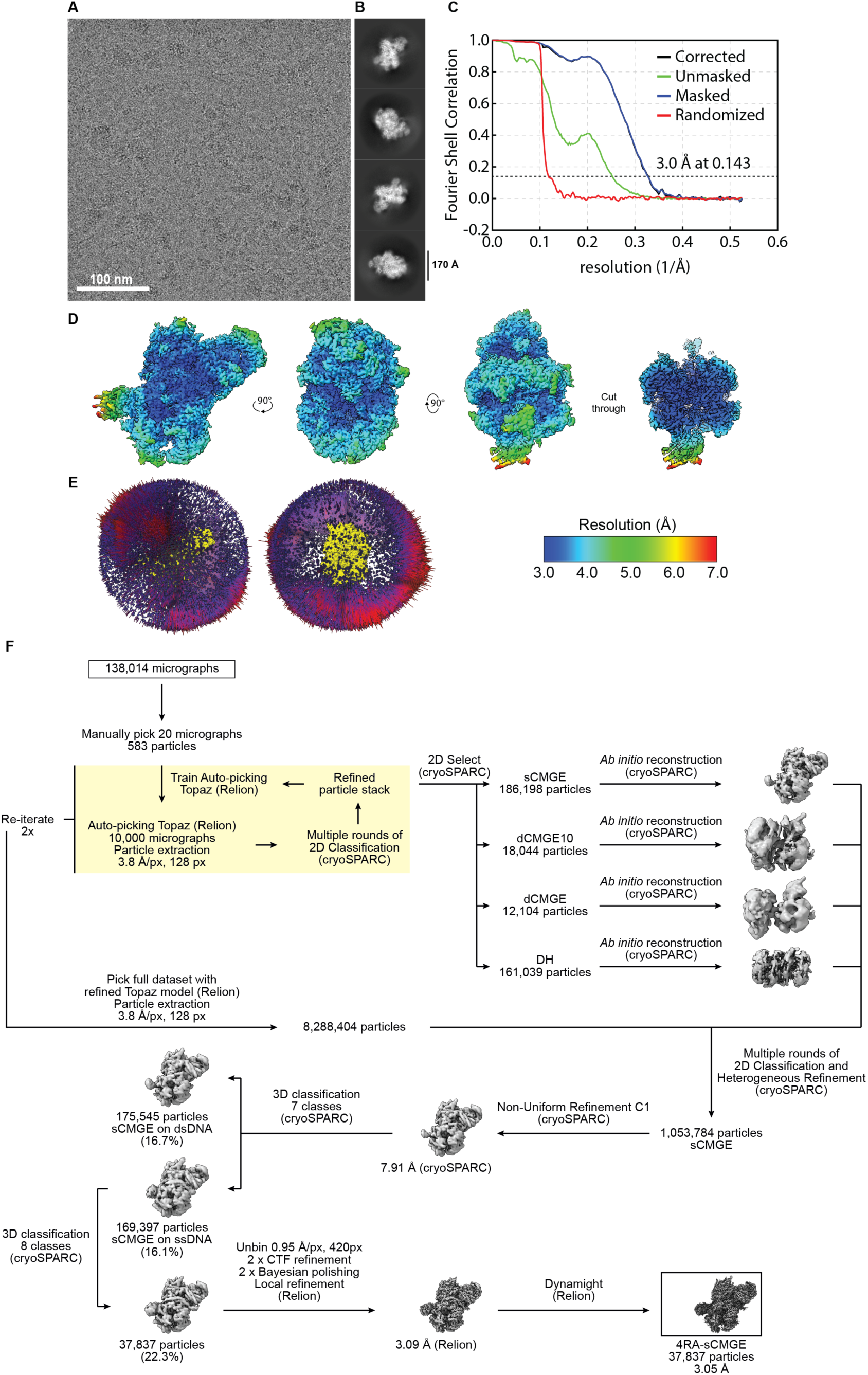
(**A**) Representative cryo-electron micrograph from the sCMGE^Mcm4RA^ dataset. (**B**) Cryo-EM 2D averages of sCMGE^Mcm4RA^. (**C**) Fourier shell correlation plot of sCMGE^Mcm4RA^. (**D**) Four views of the sCMGE^Mcm4RA^ map coloured according to the local resolution. (**E**) Angular distribution of sCMGE^Mcm4RA^. (**E**) Image processing pipeline for sCMGE^Mcm4RA^.

**Table S1.** Cryo-EM data collection, refinement and validation statistics.

|  | dCMGE10(7RA)<br>State I<br>EMDB-57126<br>PDB 29ER | dCMGE10(7RA)<br>State II<br>EMDB-57127<br>PDB 29ES | dCMGE10(7RA)<br>State III<br>EMDB-57128<br>PDB 29ET | dCMGE10(7RA)<br>State IV<br>EMDB-57130<br>PDB 29EV | sCMGE10(4RA)<br>EMDB-57131<br>PDB 29EW |
| --- | --- | --- | --- | --- | --- |
| **Data collection and processing** |  |  |  |  |  |
| Microscope | Titan Krios | Titan Krios | Titan Krios | Titan Krios | Titan Krios |
| Voltage (kV) | 300 | 300 | 300 | 300 | 300 |
| Camera | Falcon IV | Falcon IV | Falcon IV | Falcon IV | Falcon IV |
| Magnification | 130,000 x | 130,000 x | 130,000 x | 130,000 x | 130,000 x |
| Pixel size (Å) | 0.95 | 0.95 | 0.95 | 0.95 | 0.95 |
| Total electron exposure (e<sup>-</sup>/Å<sup>2</sup>) | 42.8 | 42.8 | 42.8 | 42.8 | 52.7 |
| Exposure rate (e<sup>-</sup>/px/s) | 7.1 | 7.1 | 7.1 | 7.1 | 8.74 |
| Number of frames collected | 31 | 31 | 31 | 31 | 38 |
| Defocus range (µm) | -2.0 to -2.9 | -2.0 to -2.9 | -2.0 to -2.9 | -2.0 to -2.9 | -1.6 to -2.2 |
| Automation software | EPU | EPU | EPU | EPU | EPU |
| Energy filter slit width (eV) | 10 | 10 | 10 | 10 | 10 |
| Micrographs collected | 178,563 | 178,563 | 178,563 | 178,563 | 138,014 |
| Total extracted particles | 3,553,652 | 3,553,652 | 3,553,652 | 3,553,652 | 8,288,404 |
| Final particle images | 23,793 | 67,110 | 27,138 | 76,228 | 37,837 |
| Symmetry imposed | C1 | C1 | C1 | C1 | C1 |
| Map resolution (global, Å) |  |  |  |  |  |
| FSC 0.143 (unmasked) | 7.6 | 4.5 | 7.2 | 4.5 | 3.9 |
| FSC 0.143 (masked) | 4.2 | 3.4 | 4.0 | 3.5 | 3.0 |
| **Model composition** |  |  |  |  |  |
| Non-hydrogen atoms | 108,553 | 100,889 | 100,685 | 89,838 | 56,125 |
| Protein residues | 13,224 | 12,291 | 12,261 | 10,919 | 6,859 |
| Ligands | 8/2/10/16 | 11/0/11/14 | 11/0/11/14 | 12/0/12/12 | 6/0/6/7 |
| (AGS/ADP/Mg<sup>2+</sup>/Zn<sup>2+</sup>) |  |  |  |  |  |
| DNA residues | 106 | 106 | 106 | 106 | 52 |
| **Model refinement** |  |  |  |  |  |
| Initial model used (PDB code) | 7PMK | 7PMK | 7PMK | 7PMK | 7PMK |
| Model resolution cutoff (Å) | 0.5 | 0.5 | 0.5 | 0.5 | 0.5 |
| Model-Map CC (CC<sub>mask</sub>/CC<sub>box</sub>/CC<sub>peaks</sub>/CC<sub>volume</sub>)<sup>a</sup> | 0.72/0.77/0.58/0.72 | 0.78/0.81/0.69/0.77 | 0.75/0.77/0.63/0.74 | 0.82/0.82/0.72/0.81 | 0.84/0.76/0.73/0.81 |
| Model-Map FSC 0.5 (masked/unmasked, Å)<sup>a</sup> | 4.3/4.4 | 3.7/3.7 | 4.1/4.2 | 3.7/3.7 | 3.2/3.2 |
| **Average B factors (Å<sup>2</sup>)<sup>b</sup>** |  |  |  |  |  |
| Protein | 135.87 | 118.06 | 106.87 | 116.16 | 65.72 |
| Ligand | 128.32 | 111.98 | 108.61 | 108.82 | 53.97 |
| DNA | 257.97 | 221.09 | 245.67 | 242.07 | 200.00 |
| **RMS deviations<sup>b</sup>** |  |  |  |  |  |
| Bond lengths (Å) | 0.002 | 0.002 | 0.002 | 0.003 | 0.003 |
| Bond angles (°) | 0.524 | 0.514 | 0.520 | 0.532 | 0.532 |
| **Validation<sup>b</sup>** |  |  |  |  |  |
| MolProbity score | 1.25 | 1.20 | 1.26 | 1.30 | 1.18 |
| CaBLAM outliers (%) | 0.69 | 0.89 | 0.72 | 0.87 | 1.03 |
| Clashscore | 4.69 | 4.18 | 5.02 | 3.68 | 3.73 |
| Poor rotamers (%) | 0.00 | 0.01 | 0.00 | 0.00 | 0.02 |
| C-beta deviations | 0.00 | 0.00 | 0.00 | 0.00 | 0.00 |
| **Ramachandran plot<sup>b</sup>** |  |  |  |  |  |
| Favored (%) | 97.94 | 98.27 | 98.06 | 97.27 | 97.91 |
| Allowed (%) | 2.05 | 1.70 | 1.92 | 2.72 | 2.05 |
| Disallowed (%) | 0.01 | 0.02 | 0.02 | 0.02 | 0.04 |

<sup>a</sup> Comprehensive validation (cryo-EM) in *Phenix*.
<sup>b</sup> MolProbity validation in *Phenix*.

---

**Movie S1.**

**The translocation mechanism of CMG underlies origin DNA melting.** The sequential rotary cycle of ATP binding and hydrolysis by the MCM motor that drives CMG translocation also underlies origin DNA melting by two converging CMGs. In the ATP-loaded dCMGE, MCM subunits 6, 2, 5 and 3 contact the leading strand template via their PS1 beta hairpins (“pore loops”). After transition to State I, one translocation step has been taken by Mcm7 which is now the first subunit contacting DNA at the front. We capture this state because the Mcm7 Arg finger mutation partially impairs nucleotide binding by the Mcm4 subunit, preventing it from binding DNA downstream and continuing the translocation cycle. After transition to State II, three more translocation steps have been taken and now Mcm7 is the last subunit at the back. We capture this state because the same Mcm7 Arg finger mutation also impairs ATP hydrolysis by Mcm4 thereby preventing Mcm7 from dissociating from DNA.

**Movie S2.**

**The Zinc Fingers of Mcm2 disrupt B-form DNA in the dCMGE10^Mcm7RA^.** During the transition from dCMGE to dCMGE10^Mcm7RA^ State I, the two Mcm2 Zinc Fingers come in close proximity across the double helix, creating a steric clash that destabilizes B-form DNA and drives strand separation.

**Movie S3.**

**ATPase-driven DNA shearing.** Mcm10 binding misaligns the CMGs so that a bubble is melted in the middle of the DNA. Both CMGs then hydrolyse ATP and shear the bubble further by pulling on their respective translocation strands via the MCM pore loops.

**Movie S4.**

**The mechanism of lagging strand template extraction through the Mcm2-5 interface.** In the ATP-loaded double CMGE, both DNA strands are inside the pore of the helicase. Upon transition to State I, the lagging strand template becomes entrapped between the Zinc Finger domains of Mcm2 and Mcm5. After transitioning through States II and III to State IV, the lagging strand template has been extracted out of the CMG and is fully stretched between the Zinc fingers of Mcm2 and Mcm5 such that opening of the Mcm2-5 gate would complete strand ejection. Because the stretching of the DNA is driven by ATP-hydrolysis it is irreversible, making it hard to envisage how the strand could be ejected from any gate other than Mcm2-5 in a downstream step. In conclusion, our data indicates that the lagging strand template is coming out of the Mcm2-5 gate.

